# PYM1 limits non-canonical Exon Junction Complex occupancy in a gene architecture dependent manner to tune mRNA expression

**DOI:** 10.1101/2025.03.13.643037

**Authors:** Manu Sanjeev, Lauren A. Woodward, Michael L. Schiff, Robert D. Patton, Sean Myers, Debadrita Paul, Ralf Bundschuh, Guramrit Singh

## Abstract

The Exon Junction Complex (EJC) deposited upstream of exon-exon junctions during pre-mRNA splicing in the nucleus remains stably bound to RNA to modulate mRNA fate at multiple post-transcriptional steps until its disassembly during translation. Here, we investigated two EJC disassembly mechanisms in human embryonic kidney 293 (HEK293) cells, one mediated by PYM1, a factor that can bind both the ribosome and the RBM8A/MAGOH heterodimer of the EJC core, and another by the elongating ribosome itself. We find that EJCs lacking PYM1 interaction show no defect in translation-dependent disassembly but is required for translation-independent EJC destabilization. Surprisingly, PYM1 interaction deficient EJCs are enriched on sites away from the canonical EJC binding position including on transcripts without introns or with fewer and longer exons. Acute reduction of PYM1 levels in HEK293 cells results in a modest inhibition of nonsense-mediated mRNA decay and stabilization of mRNAs that localize to endoplasmic reticulum associated TIS-granules and are characterized by fewer and longer exons. We confirmed the previously reported PYM1-flavivirus capsid protein interaction and found that human cells expressing the capsid protein or infected with flaviviruses show similar changes in gene expression as upon PYM1 depletion. Thus, PYM1 acts as an EJC specificity factor that is hijacked by flaviviruses to alter global EJC occupancy and reshape host cell mRNA regulation.

## Introduction

RNA binding proteins (RBPs) play an integral role in the synthesis and processing of RNA molecules and take center stage in their post transcriptional gene regulation(Singh et al. 2015; Hentze et al. 2018). By controlling amount, duration and location of gene expression in a cell and its various organelles, RBPs dictate proper cellular function and organismal development. Hence, it is not surprising that RBPs are mutated in several human diseases(Gebauer et al. 2021) and are targeted by pathogens to hijack cellular mechanisms for their own advantage(Girardi et al. 2021).

Among the RBP repertoire in eukaryotic cells, the Exon Junction Complex (EJC) is crucial to regulating RNA fate and is a hallmark of RNAs subjected to spliceosome-dependent intron removal(Le Hir, Saulière, and Wang 2016; Woodward et al. 2017; Schlautmann and Gehring 2020). During intron removal by the spliceosome, the three EJC core proteins EIF4A3, RBM8A (also known as Y14) and MAGOH are deposited ∼ 24 nucleotides upstream of exon-exon junctions in a sequence-independent manner(Le Hir, Moore, and Maquat 2000; Andersen et al. 2006; Bono et al. 2006). The EJC core interacts with peripheral proteins that serve as adapters to facilitate many steps in gene expression. In the nucleus, peripheral EJC factors connect it to pre-mRNA splicing (e.g., RNPS1)(Boehm et al. 2018; Schlautmann et al. 2022) and mRNA export (e.g., ALYREF)(Gromadzka et al. 2016; Viphakone et al. 2019) whereas cytoplasmic adapters link the EJC and bound RNAs to mRNA localization, translation (e.g., CASC3, also known as Barenstz or MLN51)(Palacios et al. 2004; Chazal et al. 2013; Mabin et al. 2018) and nonsense-mediated mRNA decay (NMD, e.g., UPF3)(Gehring et al. 2003, 2005; Wallmeroth et al. 2022; Yi et al. 2022). The EJC is therefore proposed to be a dynamic entity that changes composition as an mRNA proceeds through post-transcriptional events and subcellular locales. Consistently, EJCs bound to nucleus-enriched spliced RNAs contain RNPS1 and other serine-arginine rich proteins (SR proteins) while EJCs on cytoplasm-enriched spliced RNAs lack SR proteins and instead contain CASC3(Mabin et al. 2018; Singh et al. 2012; Metkar et al. 2018; Pacheco-Fiallos et al. 2023). Thus, the EJC is a key gene regulatory hub and any changes in its binding pattern or properties can alter post-transcriptional controls at multiple levels.

Human mRNAs on average have 8-9 introns (some have as many as 100) and are therefore expected to be decorated with multiple EJCs. Through formation of EJC core multimers and interactions with peripheral factors such as ALYREF and SR proteins, the EJC can package the bound RNA into compacted ribonucleoprotein (RNP) particles(Singh et al. 2012; Metkar et al. 2018; Pacheco-Fiallos et al. 2023). This so-called packaging of mRNPs mediated by EJC interactions is also consequential to post-transcriptional control. For example, EJC-mediated RNP packaging contributes to splice site selection by governing access to RNA by the spliceosome to repress cryptic 5’-splice sites(Boehm et al. 2018; Blazquez et al. 2018). Additionally, EJC-mediated mRNP formation limits m6A modification in intron-dense regions by steric inhibition of RNA modifying enzymes(He et al. 2023; Uzonyi et al. 2023; Yang et al. 2022). It remains to be seen if EJC-mediated mRNA packaging also impacts other post-transcriptional steps in gene regulation.

The last stage in the EJC life cycle is its disassembly from RNAs following which the individual subunits are imported into the nucleus for redeposition(Mingot et al. 2001; Bono et al. 2010). The elongating ribosome has been proposed as a key contributor to EJC disassembly on translated mRNAs(Dostie and Dreyfuss 2002). Indeed, a recent study estimated that a translation-dependent mechanism accounts for ∼85% of EJC disassembly in HEK293 cells(Bensaude et al. 2024). EJC removal by the translating ribosome plays a critical role in NMD as EJCs that occur in the 3’-untranslated region (3’UTR) are not subjected to such disassembly and thereby serve as a key determinant of premature translation termination(Karousis and Mühlemann 2019; Kurosaki, Popp, and Maquat 2019; Yi, Sanjeev, and Singh 2021; Carrard and Lejeune 2023). A slower and minor translation-independent EJC disassembly pathway has also been suggested(Bensaude et al. 2024), which may dismantle EJCs outside open reading frames via mechanisms yet to be ascertained.

Another activity implicated in EJC disassembly is provided by PYM1(Gehring et al. 2009). PYM1 interacts via its N-terminus with the MAGOH/RBM8A heterodimer of the EJC core and is thus aptly named Partner of Y14 and MAGOH. Interestingly, PYM1’s C-terminus shares sequence similarity with translation initiation factor EIF2A and has been shown in human cells to interact with the ribosome via its small subunit(Diem et al. 2007; Gehring et al. 2009). PYM1’s ability to associate with both the EJC and the ribosome was first proposed to mediate translation enhancement of the EJC(Diem et al. 2007). However, structural analysis of the EJC core and PYM1-MAGOH-RBM8A ternary complex suggests that the RBM8A/MAGOH heterodimer cannot simultaneously bind to PYM1 and EIF4A3, which led to a proposal that PYM1 functions in EJC disassembly(Bono et al. 2006, 2004). Indeed, PYM1 reduces EJC binding to reporter RNAs in *in-vitro* splicing assays in a concentration and MAGOH-interaction-dependent manner(Gehring et al. 2009). Further, PYM1 overexpression was found to lower EJC association with and NMD susceptibility of select spliced RNAs(Gehring et al. 2009). Consistent with an EJC disassembly function, overexpression of *Drosophila* PYM reduced EJC binding to the *oskar* mRNA and disrupted the localization of this mRNA to the oocyte posterior pole(Ghosh et al. 2014). However, in contrast to human PYM1, no ribosomal association was observed for Drosophila PYM, which can also disassemble the EJC from untranslated RNAs(Ghosh et al. 2014). Thus, it remains unresolved if PYM1 function in human cells is linked to translation-dependent EJC disassembly or if human PYM1 acts in a translation-independent manner, as does its Drosophila ortholog. Moreover, the relative contribution of PYM1 and elongating ribosome to translation-dependent EJC disassembly also remains unknown in any system.

Interestingly, human PYM1 is a target of several viruses. For example, PYM1 interaction with ORF57 protein of Kaposi’s sarcoma-associated herpesvirus enhances viral RNA translation(Boyne et al. 2010). PYM1 also interacts with capsid proteins of several viruses from the *Flaviviridae* family [e.g., West Nile virus (WNV), Zika virus (ZIKV), Dengue virus (DENV), Hepatitis C virus (HCV)](M. Li et al. 2019; Ramage et al. 2015). The flaviviral RNA genome replicates in special compartments formed from invaginations of the endoplasmic reticulum (ER)(Verhaegen and Vermeire 2024).

Interestingly, in WNV infected U2OS cells, MAGOH and RBM8A accumulate in a membrane fraction, suggesting that the EJC may be active in this stage of the flaviviral lifecycle(M. Li et al. 2019). The consequence of PYM1 sequestration by flaviviral capsid proteins in this context is unclear. Moreover, NMD suppression is a well-documented strategy employed by many viruses, presumably to promote hijacking of cellular functions and stabilize NMD-sensitive viral transcripts (reviewed in ref. (Leon and Ott 2021)). The capsid-PYM1 interaction has also been proposed to be responsible for reduced NMD activity observed in virus infected cells(M. Li et al. 2019). However, the role of PYM1, the EJC core, and NMD in viral replication or life cycle remains to be completely elucidated. To this end, it will be important to define the role of PYM1 in regulating EJC binding and function on cellular RNAs.

In this study, we investigated the role of PYM1 in regulating EJC binding to mRNA in human embryonic kidney (HEK) 293 cells. In support of PYM1 mainly contributing to a slow, translation-independent EJC disassembly mechanism, we find that disrupting the PYM1-EJC interaction does not affect translation-dependent EJC disassembly but increases EJC footprints on untranslated RNAs. Surprisingly, PYM1 interaction-deficient EJCs also accumulate on sites away from canonical EJC binding position and are enriched on longer exons and on transcripts with fewer exons. Further, acute reduction of PYM1 levels in HEK293 cells causes modest NMD inhibition.

Interestingly, PYM1 depletion has opposing effects on the stability of two classes of mRNAs that localize at or around the ER. mRNAs that localize to the ER and contain shorter and more than average number of exons are destabilized whereas mRNAs that localize to an ER-linked TIS11B-containing biomolecular condensate called TIS-granule(Ma and Mayr 2018) and are characterized by longer and fewer exons are stabilized. These two classes of mRNAs also show similar gene expression changes in human cells infected with Zika and Dengue viruses. Our results suggest that PYM1 destabilizes the EJC in a translation-independent manner to prevent EJC accumulation at non-canonical positions thus acting as an EJC specificity factor. The flavivirus capsid-PYM1 interaction potentially disrupts this activity to reshape the ER-linked host mRNA regulation.

## Results

### Translation inhibition, but not disruption of the EJC-PYM1 interaction, results in widespread changes in EJC footprints

To investigate the relative contributions of PYM1 and the elongating ribosome to EJC disassembly in human cells, we measured EJC occupancy after either blocking the EJC-PYM1 interaction or translation elongation (**Fig. 1a**). To block EJC-PYM1 interaction, we took advantage of a previously reported mutant of MAGOH (E117R, MAGOH^E117R^), which can still assemble into the EJC but does not interact with PYM1 due to mutation of a key side chain (Gehring et al., 2009, **Supplementary Fig. 1a, b**). We generated a stable HEK293 cell line expressing FLAG-tagged MAGOH^E117R^ (**Supplementary Fig. 1c**). As expected, like stably expressed wild-type (WT) FLAG-MAGOH(Singh et al. 2012), FLAG-MAGOH^E117R^ interacts with EIF4A3 and RBM8A, but unlike FLAG-MAGOH, it shows no detectable interaction with PYM1 (**Fig. 1b**). Moreover, the immunoprecipitation (IP) of FLAG-tagged wild-type and mutant MAGOH proteins shows that the FLAG-MAGOH^E117R^ EJC core does not show any change in stability under increasing salt concentrations (**Supplementary Fig. 1d**) or in its ability to co-immunoprecipitate several key EJC co-factors tested (**Supplementary Fig. 1e**). Thus, the EJC containing FLAG-MAGOH^E117R^ is likely to maintain all its known activities except for its interaction with PYM1.

**Fig. 1.**
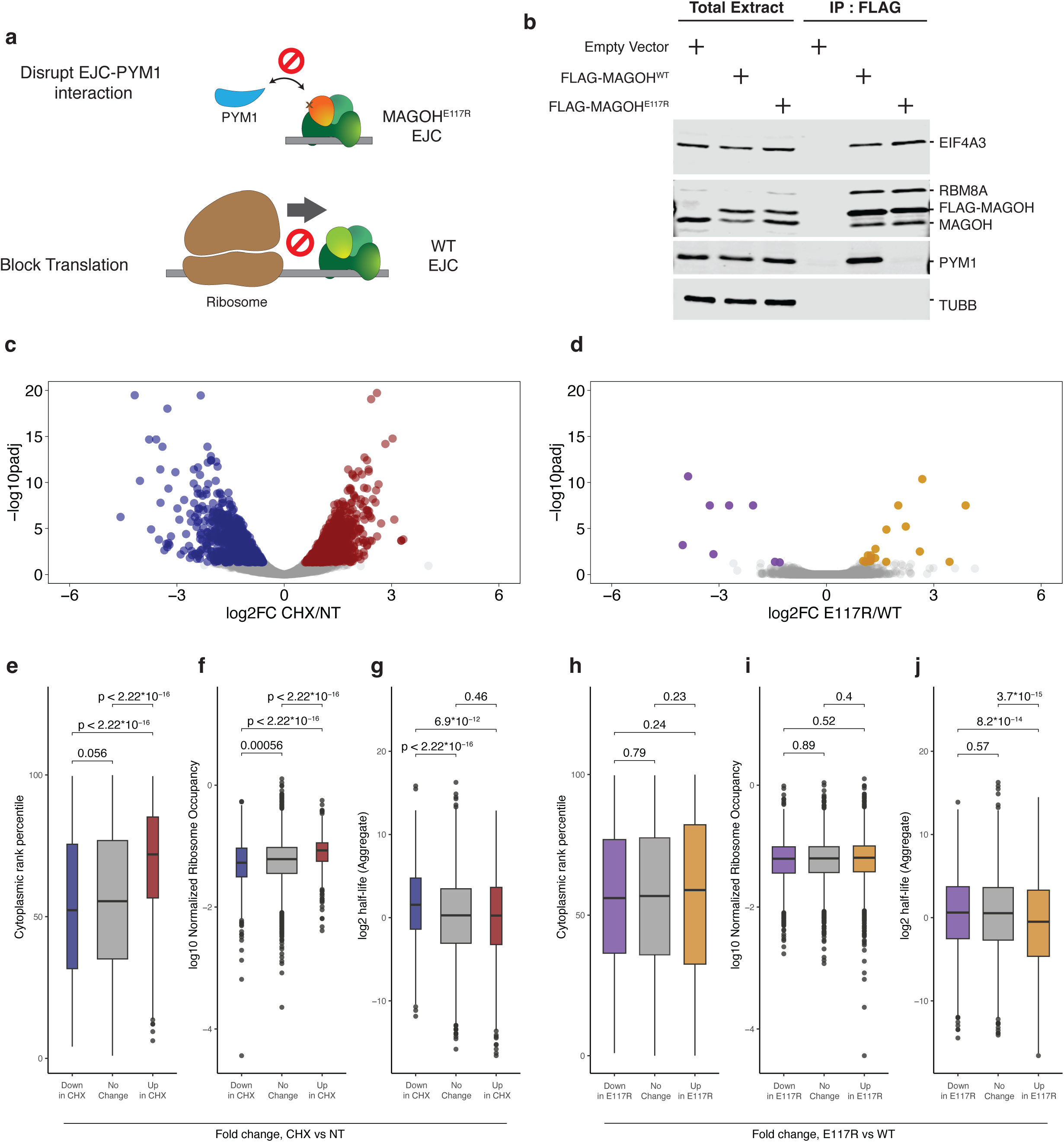
Translation but not EJC-PYM1 interaction is the major EJC disassembling activity in HEK293 cells. **a.** A schematic of the experimental approach used to disrupt EJC disassembly via disruption of EJC-PYM1 interaction using the MAGOH^E117R^ mutant (top) or using translation inhibition by cycloheximide (bottom). Gray line represents RNA and other shapes are as labeled. **b.** Immunoblots showing levels of proteins indicated on the right in total extract or FLAG immunoprecipitation (IP) fractions from cell lines indicated on top of each lane. **c, d**. Volcano plots showing the gene level changes in EJC footprints upon (**c**) CHX treatment and (**d**) MAGOH^E117R^ expression. Genes that show a significant change with *padj* < 0.05 are colored blue (downregulated) and red (upregulated). **e-j.** Boxplots comparing metrics on the y-axes among the genes with increase (up), no change or decrease (down) in EJC footprints upon CHX versus no treatment (**e**, **f**, **g**) and upon FLAG-MAGOH^E117R^ versus FLAG-MAGOH^E117R^ expression (**h**, **i**, **j**). Box plot features: box represents interquartile range (IQR), central band represents the median, whiskers extend to 1.5-times the IQR on both sides, and dots represent the outliers. *p*-values are from Wilcoxon signed-rank test comparing the two indicated distributions.

To analyze changes in EJC occupancy resulting from any defects in PYM1 mediated EJC disassembly, we performed RIPiT-Seq(Yi and Singh 2021) via tandem IP of FLAG-MAGOH (WT or E117R) and endogenous CASC3. We chose CASC3 for the second IP to enrich cytoplasmic/later-stage EJCs which are likely to be more susceptible to defects in EJC disassembly. To analyze the effect of translation elongation inhibition on EJC occupancy, both WT and the mutant MAGOH expressing cells were treated with cycloheximide (CHX) for 3 hours prior to their harvesting for RIPiT-Seq (**Supplementary Fig. 2a, b**). The two RIPiT-Seq replicates from each condition were well-correlated for gene level counts for uniquely mapping EJC footprint reads and clustered with their respective experimental condition in principal component analysis (**Supplementary Fig. 2c-g**). Therefore, individual replicates were combined for further analysis.

To quantify EJC occupancy for each protein-coding gene in the reference human transcriptome (Ensembl release 100, hg38)(Harrison et al. 2024), we estimated total EJC footprint signal across the entire length of a representative transcript (as per the MANE select annotation(Morales et al. 2022)). As expected based on a previous report that translating ribosomes remove EJCs from mRNAs(Dostie and Dreyfuss 2002), footprints of FLAG-MAGOH and CASC3 containing EJCs (MAGOH^WT^ EJC from hereon) show dramatic changes between CHX treated and untreated cells with more than 1/4^th^ of all transcripts showing significant changes in EJC footprints (**Fig. 1c**, 1,833 genes with significantly changing EJC footprints as defined by padj<0.05). In contrast, when MAGOH^WT^ EJCs were compared to those containing FLAG-MAGOH^E117R^ and CASC3 (MAGOH^E117R^ EJC from hereon), overall EJC occupancy changes significantly for only 26 genes (**Fig. 1d**). Thus, translocating ribosomes have a profound effect on EJC occupancy in HEK293 cells whereas the effect of EJC-PYM1 interaction on the EJC landscape is more subtle (also see **Supplementary Fig. 2h**). This finding agrees with a recent study by Bensaude *et al*. estimating that 85% of EJCs are disassembled via the translation machinery.

### Distinct classes of mRNAs show altered EJC occupancy upon inhibition of translation and PYM1-EJC interaction

We predicted that EJCs bound to efficiently translated, cytoplasmic mRNAs would be the most sensitive to the dramatic impact of translation elongation inhibition. Consistently, transcripts with significantly increased EJC footprints upon CHX treatment have the highest relative cytoplasmic levels (**Fig. 1e**; median cytoplasmic fraction = 0.729) whereas mRNAs that show a relative increase in EJC occupancy in untreated cells have the lowest cytoplasmic abundance (**Fig. 1e**; median cytoplasmic fraction = 0.52; *p*<2.2 × 10^-16^). Thus, localization of mRNAs from nucleus to cytoplasm promotes translation-dependent EJC disassembly. Next, transcripts with higher EJC occupancy upon CHX treatment appear to be translated more efficiently as they have a significantly higher normalized ribosome occupancy (median log_10_TE= -1.07) as compared to mRNAs with elevated EJC occupancy in untreated cells (**Fig. 1f**; median log_10_TE=-1.28; *p*<2.2 × 10^-16^). Finally, mRNAs with increased EJC occupancy upon CHX treatment have significantly shorter median half-lives (median log transformed t_½_=0.248) as compared to mRNAs enriched in EJC under normal conditions (**Fig. 1g**; median log transformed t_½_=1.555; *p*=6.9 × 10^-12^). Presumably, translation inhibition also leads to mRNA stabilization(Dave et al. 2023; Tuck et al. 2020), which in turn can contribute to the observed higher EJC signal on mRNAs upon CHX treatment. Gene Ontology (GO)-term enrichment analysis of gene sets with increased EJC footprint signal upon CHX treatment identifies keywords including nucleus, mRNA processing and mRNA splicing (**Supplementary Fig. 2i**), functional groups of genes that encode unstable mRNAs(Schwanhäusser et al. 2011). Among the set of genes comparatively enriched with EJCs in untreated cells, the most prominently enriched GO terms are for cytoplasmic ribosomes and ribosomal proteins (**Supplementary Fig. 2i**), consistent with earlier reports that translation inhibition lowers EJC occupancy on this set of mRNAs(Hauer et al. 2016; Mabin et al. 2018). Taken together, these observations affirm that cytoplasmic mRNA metabolic processes are critical in shaping EJC occupancy on mRNAs.

We next determined if disruption of EJC-PYM1 interaction influences EJC occupancy on distinct groups of mRNAs. Although FLAG-MAGOH^E117R^ EJC footprints significantly change only a few transcripts, we compared properties of genes in the top and bottom quartiles based on fold change in footprints of the mutant versus the wild-type EJC. While the EJC occupancy upon loss of EJC-PYM1 interaction shows no correlation with relative nucleocytoplasmic mRNA distribution or translation efficiency (**Fig. 1h, i**), the quartile of transcripts with the most MAGOH^E117R^ EJC footprint signal have lower half-lives as compared to those in the bottom quartile (**Fig. 1j**; median log transformed t_½_= -0.49 and 0.61 for the top and bottom quartiles, respectively; *p*=8.2 × 10^-14^). The genes with the highest mutant EJC signal are enriched in GO term keywords related to transcriptional regulation, DNA binding and ribonucleoprotein, classes that are characterized by unstable mRNAs (**Supplementary Fig. 2i**). On the other hand, genes with comparatively lower signal for the mutant EJCs (and hence more association with the wild-type EJC) are enriched in membrane related GO terms (**Supplementary Fig. 2i**). Similar membrane related GO terms are also observed in genes with reduced EJC signal upon CHX treatment. Thus, even though the disruption of EJC-PYM1 interaction generally has only a modest effect on gene-level EJC occupancy, distinct functionally related classes of genes appear to be more dependent on this interaction.

### MAGOH^E117R^ EJC shows no defect in translation-dependent disassembly but is enriched on potentially untranslated long non-coding RNAs

We next investigated the effects of EJC-PYM1 interaction on EJC occupancy in a translation-dependent and -independent manner. A direct comparison of fold changes in EJC footprints associated with translation inhibition and disruption of EJC-PYM1 interaction for each gene shows a very weak correlation between the two conditions (*R^2^*=0.12; **Fig. 2a**). Conversely, a strong correlation is observed between fold changes of the mutant and wild-type EJCs upon CHX treatment (*R^2^*=0.86; **Fig. 2b**). Thus, the loss of PYM1-MAGOH interaction does not alter the inherent ability of the EJC to disassemble during translation. Next, we hypothesized that the effect of PYM1 on EJC occupancy on long non-coding RNAs (lncRNAs) would reflect its translation-independent function. We find that, as compared to the wild-type EJC, the MAGOH^E117R^ EJC shows a significantly higher occupancy on lncRNAs (**Fig. 2c**, *p*=2.61 × 10^-7^). To rule out any potential effect of translation on lncRNA EJC occupancy, we compared the differences in MAGOH^E117R^ EJC versus MAGOH^WT^ EJC binding on the top quartile of nuclear-localized lncRNAs (based on their relative nucleocytoplasmic distribution) to all other lncRNAs. Notably, the top quartile contains lncRNAs with >95^th^ percentile rank for their nuclear localization. We observe a significantly higher MAGOH^E117R^ EJC occupancy on the nuclear lncRNAs (**Fig. 2d**, *p*=0.0263), thereby suggesting that EJC-PYM1 interaction influences EJC binding in a translation-independent manner even though it has no apparent effect on translation-dependent changes in global EJC occupancy.

**Fig. 2.**
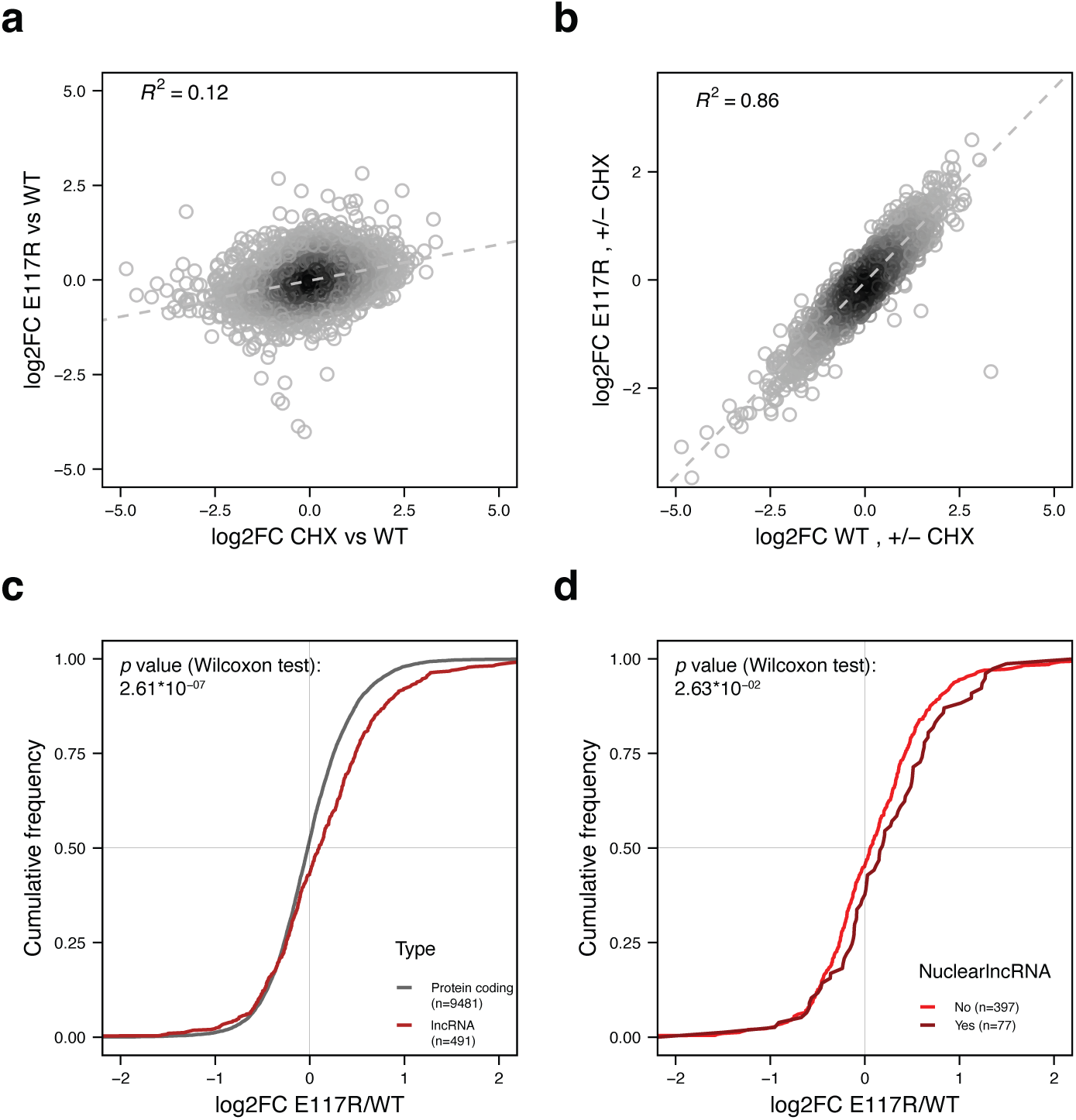
EJC-PYM1 interaction affects EJC occupancy in translation independent manner. **a.** Scatter plot comparing fold changes observed in EJC footprints on MANE select protein coding mRNAs and lncRNAs upon translation inhibition (x-axis) and disrupted EJC-PYM1 interaction (y-axis). The coefficient of determination is indicated in the top left corner. **b.** Scatter plot comparing gene-level fold changes as in (**a)** observed in CHX treated versus untreated FLAG-MAGOH^WT^ EJC footprints (x-axis) and FLAG-MAGOH^E117R^ footprints (y-axis). Only genes with baseMean>100 are included. The coefficient of determination is indicated onn the top left corner. **c.** Cumulative distribution of fold changes in footprints of FLAG-MAGOH^WT^ EJC versus FLAG-MAGOH^E117R^ EJCs on long non-coding RNAs (lncRNAs, red) as compared to protein coding genes (gray). *p*-value comparing the two distributions and numbers in each group are indicated on the top left and the bottom right, respectively. **d.** Cumulative distribution of fold changes in the mutant EJC footprints compared to wild-type as in (**c**) for two sets of lncRNAs grouped based on their nucleo-cytoplasmic ratio. The top 25% nuclear lncRNAs (nuclear rank percentile > 0.9580, dark red) are compared to all other lncRNAs (red).

### Disruption of EJC-PYM1 interaction causes EJC footprint accumulation on non-canonical locations

To investigate if the inhibition of EJC-PYM1 interaction or translation elongation affects EJC positioning on exons, EJC footprint reads from all four conditions (MAGOH^E117R^ and MAGOH^WT^ EJCs with and without CHX treatment) were plotted relative to exon-exon junctions (**Fig. 3a** and **Supplementary Fig. 3a**). Notably, in these meta-exon distributions, all genes were equally weighted, regardless of their expression levels, to prevent bias toward highly expressed genes. Also, the area under the curve for each condition remains the same so that the shape of read distribution, and hence relative EJC positioning along exons, can be compared across the conditions. As expected, EJCs from all conditions show a major peak at the canonical EJC position, ∼24 nucleotides (nt) upstream of exon 3’ ends (**Fig. 3a** and **Supplementary Fig. 3a**). Both MAGOH^WT^ and MAGOH^E117R^ EJCs show a similar footprint distribution from untreated and CHX treated cells suggesting that, even though CHX dramatically changes the overall EJC occupancy at the transcript level, the relative EJC position on exons remains unchanged. Interestingly, contrary to the expectations if PYM1 was a major disassembly factor, MAGOH^E117R^ EJC footprints are reduced at the canonical EJC position as compared to MAGOH^WT^ EJCs (**Fig. 3a**). In turn, the mutant EJC footprints show a modest increase in regions away from the -24 position (**Fig. 3a**). Previous studies have shown that EJCs can be detected at such non-canonical positions albeit at a lower level(Saulière et al. 2012; Singh et al. 2012). Our observation here suggests that the PYM1 interaction deficient EJC is more predisposed to persist in non-canonical regions than its wild-type counterpart. As we observed no differences in number of novel splice junctions detected in the RIPiT-Seq datasets from the wild-type and mutant MAGOH expressing cells (**Supplementary Fig. 3b**), the observed non-canonical EJC binding is unlikely to be due to novel or altered splicing events in the annotated exonic regions.

**Fig. 3.**
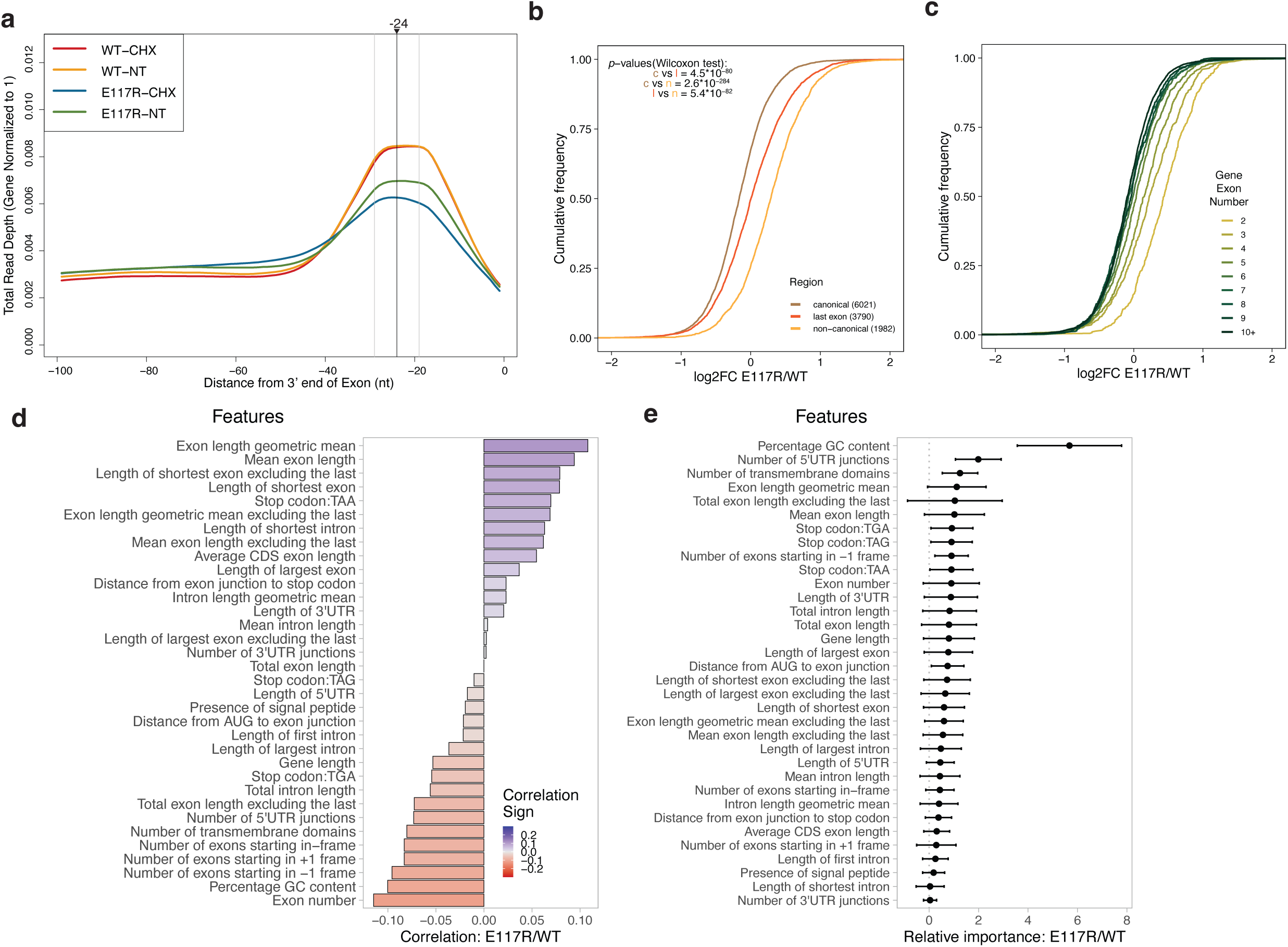
Disruption of EJC-PYM1 interaction increases EJC footprints in non-canonical regions in a gene architecture dependent manner. **a.** A meta-exon plot showing distribution of reads relative to exon 3’ ends for the four RIPiT-seq conditions shown in the top left corner. Gray lines indicate -19 and -29 nt, and the black line represents -24 nt from the 3’ end of exons. Only exons longer than 100nt in length are included. **b.** Cumulative distribution of fold changes in the mutant versus wild-type EJC footprints calculated for canonical regions, non-canonical regions and last exons within transcripts as indicated in the bottom right corner (numbers in parentheses indicate number of genes contributing to each region compared). *p*-values comparing the distributions are in the top left corner. **c.** Cumulative distribution plots as in (**b**) comparing fold changes of the mutant versus wild-type EJC footprints on genes 2 to 10+ exons, as indicated in the legend on the right. **d.** Correlation between fold-changes in the mutant versus wild-type EJC footprints and 34 different gene sequence and architectural features. The heatmap in the bottom right corner indicates color range for strength of correlation. **e.** Relative importance as calculated by machine learning analysis of shown features with gene level fold-changes in the mutant versus wild-type EJC footprints. Error bars represent one standard deviation from the calculated mean feature importance over 30 runs.

To further examine the binding of the mutant and wild-type EJCs at non-canonical positions of each spliced protein-coding MANE select transcript, we separately counted MAGOH^WT^ and MAGOH^E117R^ EJC footprint reads in (i) canonical regions, (ii) non-canonical regions of all exons with a downstream intron, and (iii) last exons. For non-canonical regions of the first to the penultimate exon, 100 nt regions from each exon’s 5’-end and 150 nt immediately upstream of exon 3’ -ends were excluded to minimize the contribution of canonical EJC footprints to non-canonical signal (see Methods for details). Also excluded from the analysis were any last exons that overlapped with known introns (based on a list compiled by Kovalak *et al*.)(Kovalak et al. 2021). Differential expression analysis between the MAGOH^WT^ and MAGOH^E117R^ EJC footprint signal in these three regions showed that as compared to the wild-type EJC the mutant EJC was most enriched in non-canonical regions (**Fig. 3b**). The mutant EJC binding was also higher in the last exons as compared to the canonical regions.

This increased propensity of the mutant EJC to bind to non-canonical regions is also supported by the enrichment of MAGOH^E117R^ EJC over the wild-type EJC on genes with fewer than average number of exons (i.e., lower intron density and hence longer non-canonical regions). Strikingly, we observe a progressive increase in mutant over wild-type EJC enrichment as exon count in genes decreases, particularly from five to two (**Fig. 3c**). These observations suggest that the loss of PYM1 interaction allows the EJC to persist longer or bind more often to non-canonical locations within long exonic stretches of RNA, thereby reducing the EJC binding specificity. Importantly, translation elongation inhibition does not cause such an increase in non-canonical EJC occupancy (**Supplementary Fig. 3c, d**).

Motivated by the above findings, we systematically examined the relationship between PYM1’s influence on EJC occupancy and 34 different gene sequence and architectural features (**Fig. 3d**). We find that the top two features directly correlated with higher mutant EJC binding pertain to exon length (e.g., exon length geometric mean has Kendall’s τ =0.108, *p*=1.35 × 10^-50^) (**Fig. 3d**). On the other hand, the top two features anti-correlated to the preferred mutant EJC binding are exon number (τ = - 0.115, *p*=2.02 × 10^-54^) and GC content (τ =-0.1, *p*=9.01 × 10^-44^) (**Fig. 3d**). Next, we took an unbiased machine learning based approach to assess the relative importance of the 34 gene features in predicting changes in mutant versus wild-type EJC binding. We permuted the values of one feature at a time and calculated the loss in the model’s ability to predict fold change in mutant versus wild-type EJC binding (see Methods for details). This analysis suggested that GC content is the most important feature for accurate prediction of the change in mutant EJC binding (**Fig. 3e**). Features pertaining to exon length and number are also observed among the top ten features in this analysis although their contribution to the fold change prediction is not significant. It is noteworthy that GC content was previously shown to be negatively correlated with exon number(Mordstein et al. 2020), which underscores their individual correlation to mutant EJC binding predicted by our model. Overall, these results confirm that disruption of PYM1-EJC interaction increases EJC occupancy on genes with fewer but longer exons that also have a higher GC content. While GC content also comes out as an even better predictor of increased EJC binding upon CHX treatment (**Supplementary Fig. 4**), both the mutant and wild-type EJC show similar relationships to various gene features in this condition, consistent with our earlier results that the loss of PYM1 binding does not affect translation-dependent EJC disassembly.

### Non-canonical EJC binding is observed on unspliced RNAs and is further enhanced for MAGOH^E117R^ EJC

By definition, any EJC binding on unspliced RNAs will be considered non-canonical binding. Although the EJC is not expected to be deposited on RNAs transcribed from intronless genes, which are not acted upon by the spliceosome, our observation of increased MAGOH^E117R^ EJC binding in other intron lacking RNA segments like 3’UTRs prompted us to test for occupancy of the mutant and wild-type EJCs on single exon mRNAs. Importantly, we filtered single-exon genes to limit our analysis to only those that do not contain any known introns(Kovalak et al. 2021). Strikingly, when single exon mRNAs are compared as a group to multi-exon mRNAs, a highly significant enrichment of MAGOH^E117R^ over MAGOH^WT^ EJC footprints is observed on these unspliced mRNAs (**Fig. 4a**). Moreover, MAGOH^E117R^ EJC enrichment on single exon mRNAs is even more pronounced than spliced mRNAs containing two exons, which show the highest mutant EJC enrichment among the spliced mRNAs (**Fig. 4a**, inset). When MAGOH^E117R^ and MAGOH^WT^ EJC footprint counts on single exon mRNAs are compared to other non-canonical regions (3’UTRs and sites away from -24 position in spliced exons), the highest enrichment of the mutant EJC footprints is again observed on single exon mRNAs (**Fig. 4b**). These findings suggest that under normal conditions EJC-PYM1 interaction limits non-canonical EJC binding, including on unspliced mRNAs.

**Fig. 4.**
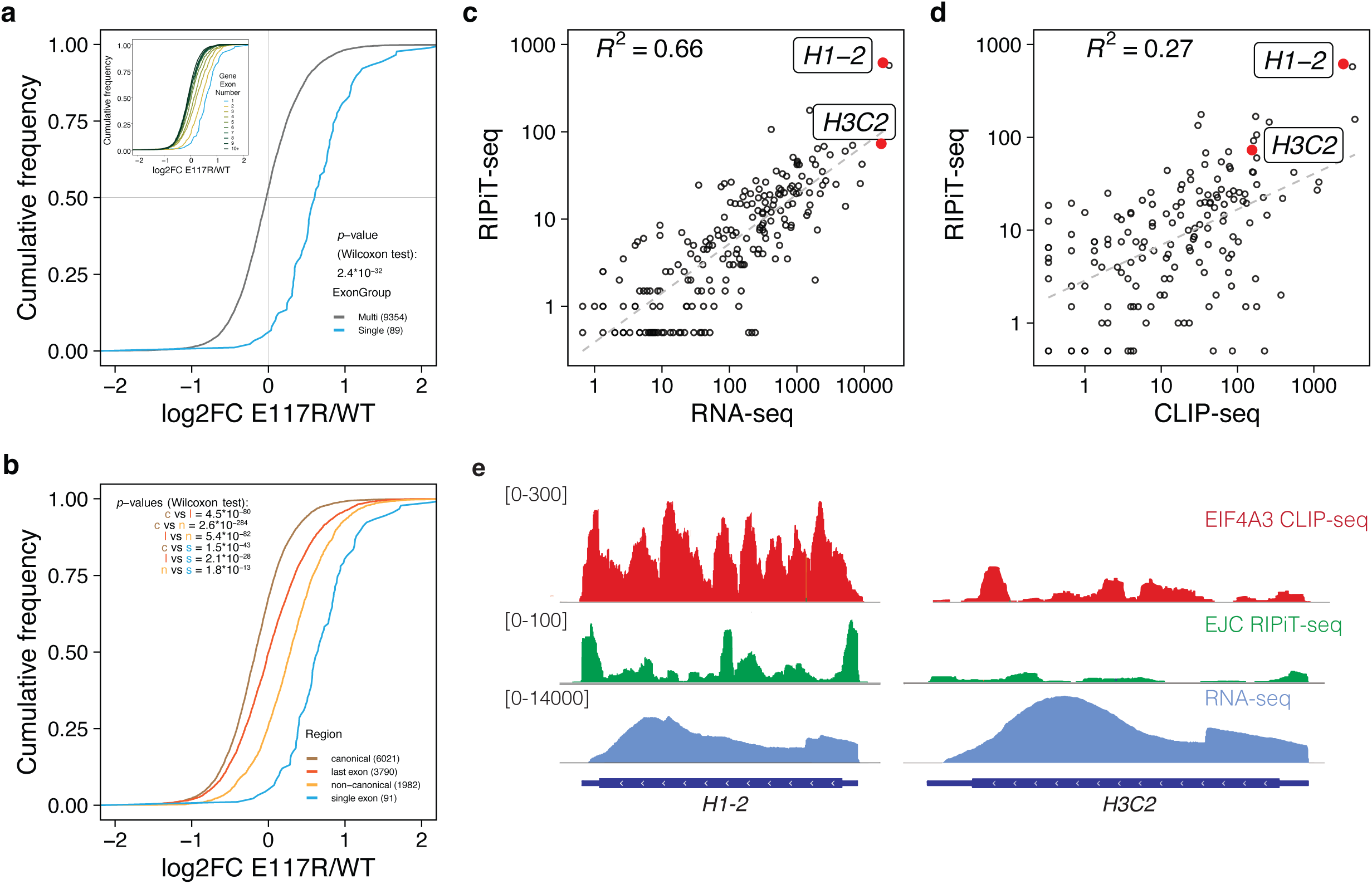
Footprints of PYM1 interaction deficient EJC are enhanced on intronless transcripts. **a.** Cumulative distribution plot showing fold changes in FLAG-MAGOH^WT^ EJC versus FLAG-MAGOH^E117R^ EJC footprints on multi-exon and single exon protein coding transcripts, as indicated in the bottom right corner. Wilcoxon test p-value is also shown. Inset shows the same plot with multi exon genes grouped based on exon number as indicated in the legend in the bottom right corner. **b.** Cumulative distribution plot as in (**a**) showing fold changes in EJC footprints calculated for single exon transcripts as compared to canonical regions, non-canonical regions and last exons from multi-exon transcripts. **c.** Scatter plot comparing signal for single exon protein-coding transcripts in FLAG-MAGOH^WT^ EJC RIPiT-seq (y-axis) and total RNA-seq (x-axis) in HEK293 cells. Gray dashed line shows best fit line with coefficient of determination in the top left corner. Two histone mRNAs shown in (**e**) are labeled. **d.** Scatter plot as in (**c**) comparing signal on single exon protein coding transcripts in FLAG-MAGOH^WT^ EJC RIPiT-seq (y-axis) and EIF4A3 CLIP-seq (ref. (Hauer et al. 2016); x-axis). **e.** Read coverage from EIF4A3 CLIP-seq, FLAG-MAGOH^WT^ RIPiT-seq and input RNA-seq on *H3C2* and *H1-2* gene loci. Numbers above each track on the left are the y-axis ranges applicable to both the gene loci.

We next sought to further characterize the unexpected EJC binding on single-exon mRNAs. At least 127 single exon mRNAs that lack any evidence of splicing show MAGOH^WT^ EJC footprint signal in HEK293 cells, suggesting that EJC binding to unspliced transcripts is not uncommon. As expected, EJC footprint detection on these mRNAs is correlated with their RNA expression levels (*R^2^* = 0.66, **Fig. 4c**). Interestingly, single exon genes with similar RNA expression levels can exhibit large differences in EJC footprints as exemplified by two histone protein encoding genes, *H1-2* and *H3C2* (**Fig. 4c**). EJC binding in non-canonical regions, including on unspliced mRNAs in RIPiT-seq datasets, could be due to an artificial assembly of EJC-like complexes on RNA post cell lysis(Mili and Steitz 2004). To test whether the non-canonical EJC binding on unspliced mRNAs can be observed *in situ*, we analyzed a published CLIP-Seq dataset that captures RNA segments directly UV-crosslinked to EIF4A3, the RNA binding subunit of the EJC(Hauer et al. 2016). As UV-crosslinking of the RNA-protein interaction surface happens prior to cell lysis, CLIP-seq crosslinking sites provide more concrete evidence for RNA-protein interaction inside cells. We find that EJC RIPiT-seq read counts from HEK293 cells and EIF4A3 CLIP-seq read counts from HeLa cells are correlated on single exon mRNAs (**Fig. 4d**, *R^2^* = 0.27). Importantly, the EIF4A3 CLIP-seq read distribution on *H1-2* and *H3C2* genes exhibits a similar trend (**Fig. 4e**) with *H1-2* showing a much higher coverage of CLIP-seq reads as compared to *H3C2* despite similar expression level of the two genes in HeLa cells. These observations further strengthen the evidence for EJC binding to unspliced RNAs. Although mechanisms that may dictate binding and specificity of non-canonical EJC on unspliced mRNAs remain unknown, it is possible that splicing-independent association of EJC subunits to active transcription regions could contribute to such EJC binding(Choudhury et al. 2016; Akhtar et al. 2019).

### PYM1 depletion mildly upregulates endogenous NMD targets

We next sought to investigate the role of PYM1 in maintaining gene expression in HEK293 cells. A 72-hour siRNA-treatment was used to deplete PYM1 to nearly a fifth of its normal levels in this cell line (**Supplementary Fig. 5a**). Such PYM1 depletion did not alter EJC-dependent splicing events (**Supplementary Fig. 5b**) or global translation as measured by puromycin incorporation into nascent polypeptides (**Supplementary Fig. 5c, d**). Therefore, we next considered the effect of PYM1 depletion on NMD, which was previously shown to be influenced by PYM1 using NMD reporter RNAs(Gehring et al. 2009). We performed total RNA-Seq after ribosomal RNA depletion from the knockdown and control cells and quantified gene expression at the transcript level. To assess the effect of PYM1 on NMD of endogenous mRNAs, we focused on transcripts that contain premature termination codons (PTC), which are translation stops that occur >50 nucleotides from the downstream exon junction. As a control for the effect of PYM1 knockdown on other processes (e.g., transcription), these PTC+ transcripts were compared to other isoforms from the same genes that do not contain a PTC (PTC-).

When we consider all PTC+ transcripts detected in HEK293 cells, a modest but significantly higher fold change is observed for PTC+ transcripts as compared to their PTC-counterparts (**Supplementary Fig. 5e**). When we limited the PTC+ group to transcripts that are > 1.5-fold upregulated in HEK293 cells double-depleted of SMG6 and SMG7(Boehm et al. 2021) and therefore most susceptible to NMD, the effect of PYM1 depletion on this subset of PTC+ transcripts is more pronounced (**Fig. 5a**). Still, the global NMD defect upon PYM1 depletion (median log2FC of PTC+ transcripts = 0.05) is not as strong as that observed upon depletion of CASC3 (median log2FC of PTC+ transcripts = 0.38)(Gerbracht et al. 2020) or SMG6 and SMG7 co-depletion (median log2FC of PTC+ transcripts = 0.65)(Boehm et al. 2021) in HEK293 cells (**Supplementary Fig. 5f**). Among the select NMD targeted transcripts tested by quantitative RT-PCR based assay, only SRSF3 PTC+ mRNA shows a significant upregulation upon PYM1 knockdown whereas all tested NMD targets are robustly upregulated upon EIF4A3 depletion (**Fig. 5b**). Thus, NMD efficiency in PYM1 knockdown cells is reduced, but only modestly. Presumably, PYM1 deficiency leads to longer EJC lifetimes causing the complex to accumulate within untranslated RNPs and non-canonical locations thereby reducing canonical EJC deposition on newly spliced RNAs, and hence a minor NMD defect.

**Fig. 5.**
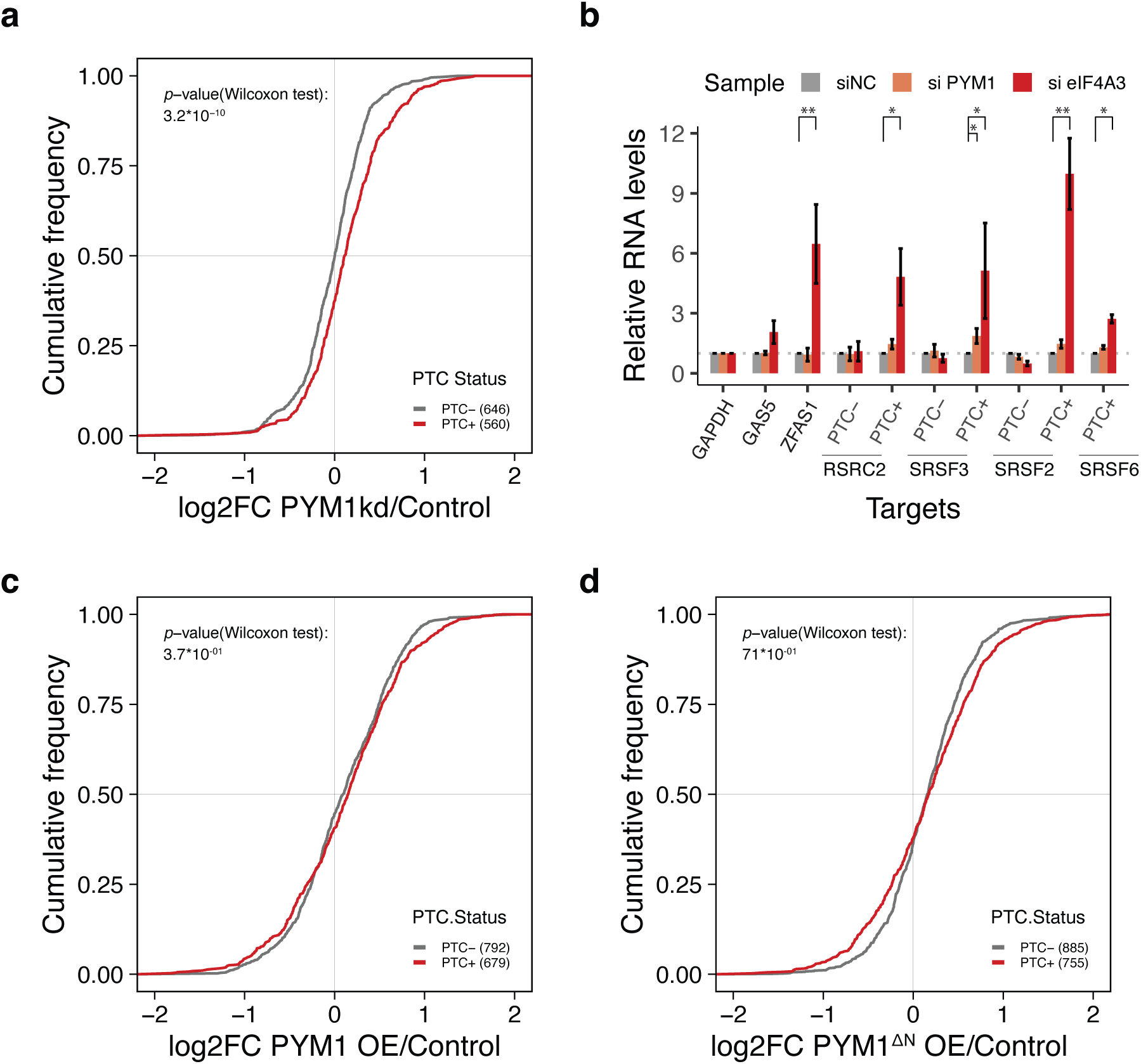
Both PYM1 depletion and overexpression lead to weak NMD suppression. **a.** Cumulative distribution of fold-changes in abundance of PTC-containing transcripts (PTC+, red line) and isoforms without a PTC from the same genes (PTC-, gray line) upon PYM1 depletion. Only those PTC transcripts that are sensitive to SMG6+7 depletion were included (see Methods). PTC status and number of transcripts in each set are indicated on the bottom right, and Wilcoxon test *p*-value is indicated on the top left. **b.** Quantitative RT-PCR based estimation of changes in expression of select NMD targeted transcripts (PTC+) and their control non-NMD isoforms (PTC-) upon transfection of control siRNA (siNC, gray bars) PYM1 siRNA (orange bars) and EIF4A3 siRNA (red bars). In each knockdown, transcript levels were first normalized to GAPDH and then fold-change in transcript levels upon PYM1 or EIF4A3 knockdown were calculated as compared to the control knockdown. Bar height in knockdown samples represents mean of 3 biological replicates and error bars indicate standard error of means. Asterisks represent *p*-values from Student’s single-tailed t-test (* < 0.05; ** < 0.01). **c**, **d**. Cumulative distribution plots as in (**a**), for PYM1 overexpression (**c**) and PYM1^ΔN^ overexpression (**d**).

A previous study suggested that PYM1 overexpression can inhibit NMD of PTC-containing TCRβ and β-globin reporters by promoting EJC disassembly(Gehring et al. 2009). To test if elevated PYM1 levels can cause a similar upregulation of endogenous NMD targets, we stably expressed in HEK293 cells N-terminal HA-tagged full-length PYM1 or its EJC interaction domain lacking deletion mutant, PYM1^ΔΝ^ (∼5-fold and ∼3-fold overexpression, respectively, **Supplementary Fig. 5g**). We confirmed that the deletion of N-terminal of PYM1 disrupts the interaction of PYM1 with MAGOH (**Supplementary Fig. 5h**). However, the exogenous expression of full-length PYM1, like that of control PYM1^ΔΝ^, did not cause any upregulation of SMG6 and 7 sensitive PTC+ transcripts (**Fig. 5c, d**) or all expressed PTC+ transcripts (**Supplementary Fig. 5i, j**).

Thus, even ∼5-fold increase in PYM1 levels does not appear to cause sufficient EJC destabilization to alter NMD of endogenous PTC+ mRNAs in HEK293 cells.

### PYM1 depletion has a differential effect on the stability of mRNAs localized to ER-linked compartments

It is possible that reduced EJC specificity in PYM1 depleted cells also affects gene expression beyond NMD. To investigate this further, GO term enrichment analysis showed that transcripts up-and downregulated upon PYM1 depletion belong to distinct functional classes (**Supplementary Fig. 6a**). Strikingly, the top five GO terms enriched among the downregulated transcripts are related to membranes and membrane-linked processes (**Supplementary Fig. 6a**). The specific downregulation of membrane targeted transcripts in response to PYM1 depletion is also observed upon independent analysis of mRNAs translated at the ER (definition based on Jan et al., 2014(Jan, Williams, and Weissman 2014)), those encoding ER-targeting sequences (signal peptide or transmembrane domain), and the ones targeted to mitochondria and peroxisomes (**Supplementary Fig. 7a**).

Interestingly, a recent study has suggested a link between mRNA expression at the ER and gene architectural features such as exon number and exon length(Ma and Mayr 2018; Horste et al. 2023). mRNAs enriched in the ER were found to have higher than average exon count. On the other hand, mRNAs localized within an ER-linked membrane-less condensate known as TIS-granule (TG) have longer and fewer exons. These findings have intriguing parallels with our observations that membrane targeted mRNAs are downregulated upon PYM1 knockdown and those with long exons are enriched in footprints of PYM1 interaction-deficient EJCs (**Fig. 3d**). Thus, PYM1 (and hence EJC) function could be linked to gene architecture and mRNA expression. Indeed, we find that in PYM1 depleted HEK293 cells, mRNAs preferentially localized to TGs and the ER are significantly up-and downregulated, respectively (**Fig. 6a, b**). When we compare these two classes of mRNAs for binding to wild-type versus the PYM1-interaction deficient EJCs, we observe a similar although weaker trend (**Fig. 6c**). Notably, the increased binding of the mutant EJC on TG+ mRNAs is mainly in the non-canonical regions (**Fig. 6d**).

**Fig. 6.**
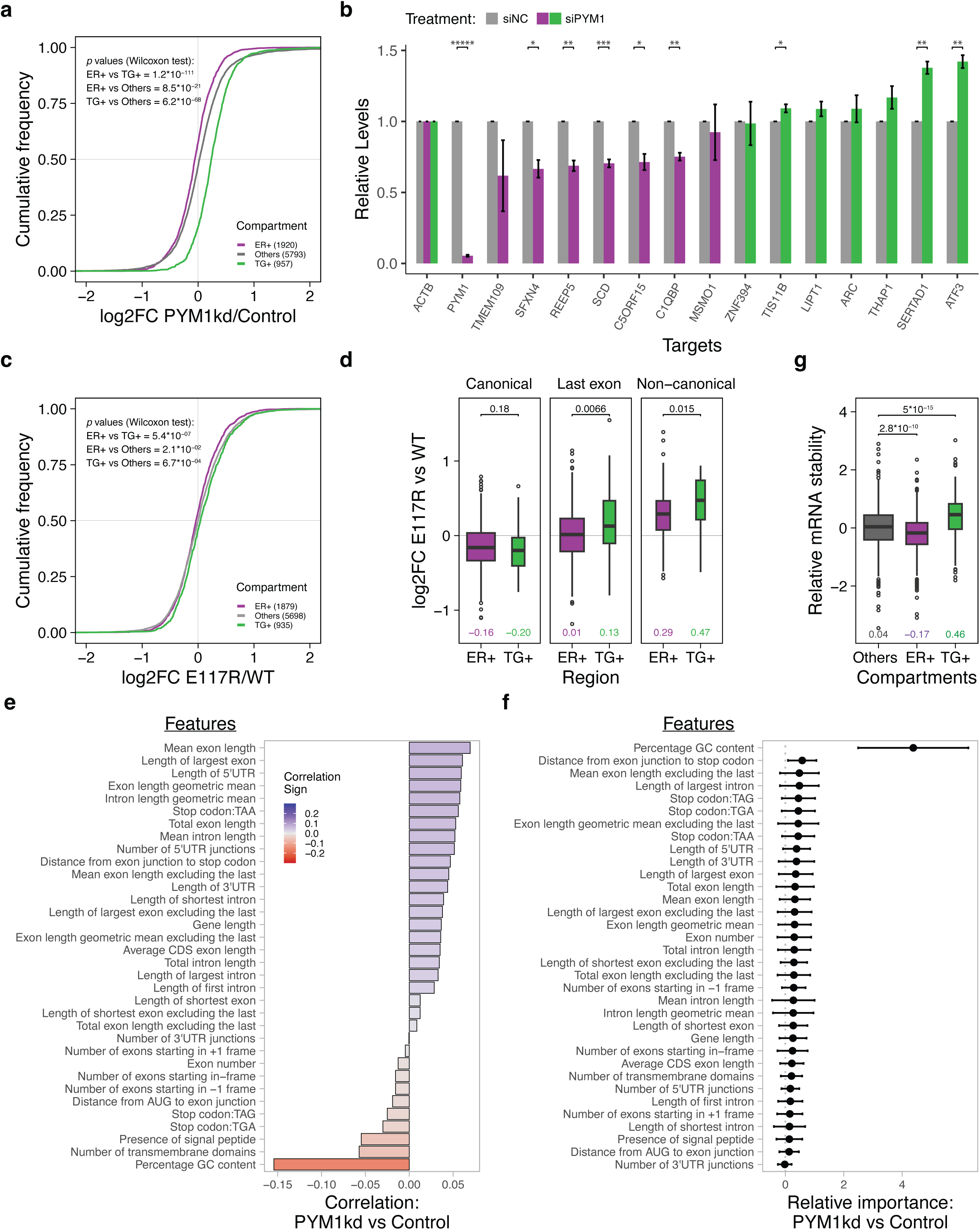
PYM1 depletion post-transcriptionally alters expression of specific classes of ER-associated mRNAs with distinct architectural features. **a.** Cumulative distribution of fold-changes upon PYM1 depletion in levels of transcripts that co-fractionate with ER (purple) and Tis-granule (green) markers as compared to other transcripts (gray); transcript classification is based on Horste et al.; numbers in parentheses indicate transcript count in each category. *p*-values from Wilcoxon test comparing the distributions are in the top left corner. **b.** Quantitative RT-PCR based validation of changes in ER+ (purple bars) and TG+ transcripts (green bars) upon PYM1 knockdown (siPYM1) as compared to negative control knockdown (siNC). Data normalization is as in Fig. 5b. Bar height in siPYM1 samples represents mean of 3 biological replicates and error bars indicate standard error of means. Asterisks represent p-values from single-tailed Student’s t-test (* < 0.05; ** < 0.01; ***** < 0.00001). **c.** Cumulative distribution of fold-changes in footprints of EJC containing FLAG-MAGOH^E117R^ as compared to FLAG-MAGOH^WT^ on ER+ (purple), TG+ (green) and other mRNAs (gray) as in (**a**). **d.** Boxplots showing fold-changes in FLAG-MAGOH^E117R^ as compared to FLAG-MAGOH^WT^ EJC footprints on canonical (left), last exon (middle) and non-canonical (right) RNA regions as indicated on top for TG+ and ER+ mRNAs. The numbers below each box plot indicate medians for each plot. Wilcoxon test *p*-values comparing the indicated distributions are above the box plots. **e.** Correlation between fold-changes in transcript expression upon PYM1 knockdown and 34 gene sequence/architecture features listed. The heatmap in the top left corner indicates color range for strength of correlation. **f.** Relative importance 34 gene sequence/architecture features with gene level fold-changes in PYM1 versus control knockdown as determined by machine learning analysis. Error bars represent one standard deviation from the calculated mean feature importance over 30 runs. **g.** Box plots comparing estimated relative stabilities of transcripts from ER+, TG+ and Other gene categories. Wilcoxon test *p*-values (top) and medians (bottom) are as indicated.

The relationship between PYM1 function and gene architecture is further confirmed by direct correlations between changes in mRNA expression upon PYM1 knockdown and exon length-based features (**Fig. 6e**). Conversely, GC content, which is intimately linked to lower exon count(Mordstein et al. 2020), is the most anti-correlated feature to altered gene expression in PYM1 depleted cells (**Fig. 6e**). Of all the features tested in our machine learning based analysis, high GC content is also the most important predictor of the observed changes in gene expression upon PYM1 depletion (**Fig. 6f**), as it was for the increased binding to PYM1-deficient EJCs (**Fig. 3d**). In the PYM1 depleted cells, a small but significant downregulation of mRNAs in the top quartile based on exon number as compared to those in the bottom quartile remains when limiting to only non-ER+ transcripts (**Supplementary Fig. 7b**). Thus, the link between PYM1 function and gene architecture likely exists independently of ER/membrane localization.

We next sought to determine if the change in ER+ and TG+ mRNA levels upon PYM1 knockdown is due to transcriptional or post-transcriptional mechanism(s). To this end, an estimation of the change in stability of these classes of mRNAs showed that as compared to other mRNAs, ER+ mRNAs are destabilized whereas TG+ mRNAs are stabilized (**Fig. 6g**, median relative stability: ER+ = -0.17, TG+ = +0.46, Others = +0.04; *p*=2.8 × 10^-10^: ER+ vs Others, *p*=5.0 × 10^-15^: TG+ vs Others). This destabilization of ER+ mRNAs is possibly caused by a mechanism that is unrelated to general NMD (**Supplementary Fig. 7c**), the NBAS-dependent ER-linked NMD (**Supplementary Fig. 7d, g**)(Longman et al. 2020), the ER-linked mRNA downregulatory activity of UPF1 long-loop isoform (**Supplementary Fig. 7e, g**)(Fritz et al. 2022), or thapsigargin-induced ER stress response (**Supplementary Fig. 7f**)(Fritz et al. 2022).

### Effects of PYM1 knockdown on ER-linked mRNA expression and NMD are recapitulated in flavivirus infected human cells

Recent biochemical screens for host proteins that interact with flavivirus capsid proteins have identified PYM1 as one of the key targets(Coyaud et al. 2018; Ramage et al. 2015; M. Li et al. 2019). This flavivirus capsid-PYM1 interaction reportedly aids in viral replication and causes an enrichment of EJC core proteins in a membrane fraction(M. Li et al. 2019). Notably, flaviviruses remodel ER membranes to create special compartments where the viral RNA genome is replicated. Our observation that PYM1 knockdown alters stability of ER and TG associated mRNAs raises a hypothesis that, upon infection, the flavivirus capsids sequester free PYM1 molecules to reshape ER-linked mRNA expression, as observed upon PYM1 depletion. To test this prediction, we first confirmed that 2X Strep-tagged WNV and DENV capsid proteins co-immunoprecipitate with FLAG-tagged PYM1 (**Fig. 7a** and **Supplementary Fig. 8a**). In support of our hypothesis, transient WNV or DENV capsid protein expression in HEK293 cells led to upregulation of TG+ transcripts and downregulation of ER+ mRNAs (**Fig. 7b** and **Supplementary Fig. 8b-d**). Notably, these trends are similar to although weaker than those observed in PYM1 depleted cells (**Fig. 6a**). Interestingly, capsid protein expression also caused elevated levels of PTC+ mRNAs as compared to their PTC-counterparts (**Fig. 7c** and **Supplementary Fig. 8e**). To examine if flavivirus infection also leads to similar changes in expression of TG+ and ER+ mRNAs, we compared expression of these groups of mRNAs in the available RNA-seq datasets from human hepatoma cells (Huh7) infected with ZIKV or DENV(Ooi et al. 2019) and human neuronal stem cells (hNSC) infected with ZIKV(Dang et al. 2019). In agreement with our experiments with transient capsid expression, all three flavivirus-infected samples displayed significant and robust upregulation of TG+ mRNAs, whereas ER+ mRNAs are weakly repressed (**Fig. 7d, e** and **Supplementary Fig. 8f**). The infected cells also exhibit reduced NMD activity as indicated by significantly increased levels of PTC+ as compared to PTC-transcripts, which is consistent with the transient capsid expression experiments (**Fig. 7f, g** and **Supplementary Fig. 8g**). Overall, these findings suggest that in flavivirus infected cells, the viral capsid protein-PYM1 interaction may limit PYM1 function in regulating EJC occupancy to thereby alter gene expression in ER-associated compartments to aid viral replication.

**Fig. 7.**
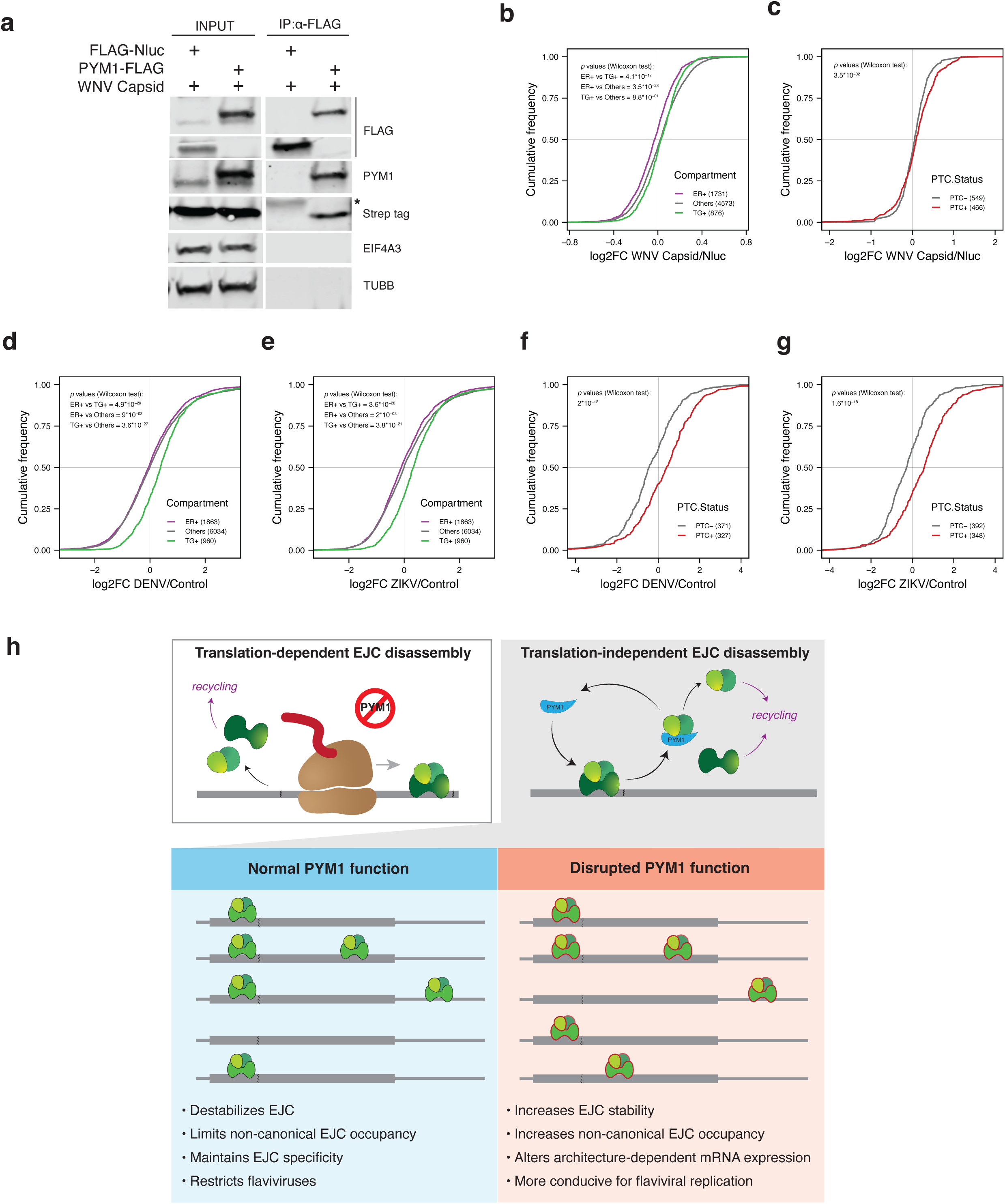
Gene expression changes upon flavivirus capsid expression or upon flaviviral infection resembles alterations observed upon PYM1 depletion. **a.** Western blot showing levels of proteins on the right in input and FLAG-IP fractions of cells expressing proteins indicated above each lane. **b**, **d**, **e**. Cumulative distribution of fold-changes of ER+, TG+ and Other category of transcripts upon WNV capsid expression in HEK293 cells (**b**), DENV infected Huh7.5.1 cells (Ooi et al.) (**d**), and ZIKV infected Huh7.5.1 cells (Ooi et al.) (**e**). Wilcoxon test *p*-values comparing the distributions (top left corner) and a legend with transcript counts in the three categories (bottom right) are shown. **c**, **f**, **g**. Cumulative distribution of fold-changes in transcript abundance of PTC-containing transcripts (red line) and isoforms without a PTC from the same genes (gray line) upon expression of WNV capsid in HEK293 cells (**c**), DENV infected Huh7.5.1 cells (Ooi et al.) (**f**), and ZIKV infected Huh7.5.1 cells (Ooi et al.) (**g**). Wilcoxon test *p*-values comparing the two distributions (top left corner) and a legend with transcript counts in the two categories (bottom right) are shown. **h.** Model for EJC disassembly in human cells and contribution of PYM1 to EJC binding specificity. Top: Two independent modes of EJC disassembly. Left: Disassembly of EJC (green shapes) from mRNA (gray line) by the ribosome (brown) does not require PYM1; jagged line on mRNA represents splice-junction. Disassembled EJC subunits are recycled and re-deposited. Right: PYM1 (blue) mediates translation-independent EJC subunit removal from RNA for subsequent sub-unit recycling. Bottom: Multiple molecules of a hypothetical spliced RNA depicting steady-state EJC occupancy under two conditions. Left: Under normal conditions, PYM1 limits EJC occupancy in non-canonical regions to maintain higher EJC binding specificity upstream of exon junctions (jagged line). Right: When PYM1 function is disrupted (e.g., in flavivirus infected cells), EJCs become more stable (indicated by red outline of shapes denoting EJC subunits) that causes the complex to persist longer in untranslated regions and RNAs transcriptome-wide (not shown). This leads to reduced EJC subunit recycling, a lower new canonical EJC deposition, and hence an overall decrease in EJC specificity.

## Discussion

### PYM1 destabilizes the EJC in a translation-independent manner to maintain canonical EJC occupancy

By comparing the two activities implicated in EJC disassembly, we provide evidence that in human cells the translating ribosome is a major contributor to EJC disassembly as compared to PYM1 (**Fig. 1**). Moreover, PYM1 is not required for the complex removal during translation and instead acts in a translation-independent manner to destabilize the EJC (**Fig. 2** and **Fig. 7h**). Our results are also consistent with recent estimates that at steady state a major fraction of EJCs are disassembled in a translation-dependent manner(Bensaude et al. 2024). Further, the slower and minor translation-independent EJC disassembly pathway described by Bensaude *et al*. is likely to be mediated by PYM1. Such a function for PYM1 in translation-independent EJC disassembly is also consistent with the observations from Drosophila embryos where PYM overexpression reduces EJC occupancy on a reporter RNA that does not undergo translation(Ghosh et al. 2014). Even though we measured PYM1’s translation-independent effect on EJC stability only on lncRNAs, it is likely that PYM1 affects EJC occupancy on translated RNAs before they engage with ribosomes. Such a possibility is supported by a minor but noticeable effect on the occupancy of PYM1-interaction deficient EJC on protein-coding RNAs (**Fig. 1** and **2**). Thus, PYM1 can act globally on all RNPs independently of translation to destabilize EJCs.

A striking change in occupancy of the PYM1-interaction deficient EJCs is their accumulation away from the canonical EJC binding sites. This increased non-canonical MAGOH^E117R^-EJC binding is observed in long exonic regions of spliced RNAs, (**Fig. 3**) and, most surprisingly, on unspliced mRNAs (**Fig. 4**). EJC binding away from exon junctions in human cells has been documented in multiple studies, even in those that relied on more stringent in situ EIF4A3-RNA crosslinking using ultraviolet light(Hauer et al. 2016; Saulière et al. 2012). We show that non-canonical binding of WT EJCs on unspliced RNAs can be detected via both crosslinking-dependent and -independent approaches (**Fig. 4**), thus confirming it to be a *bona fide* EJC property. How the EJC gets to non-canonical locations remains unknown. It is possible that such binding patterns could be due to the complex sliding on RNA after splicing-dependent deposition. Alternatively, mechanisms that are independent of splicing, e.g., the engagement of EJC factors with the transcriptional machinery(Akhtar et al. 2019) or nascent transcripts(Choudhury et al. 2016) could recruit EJC subunits to assemble in non-canonical locations on spliced as well as unspliced RNAs. While mechanism(s) of origin of non-canonical EJCs remain an important open question, our findings suggest that a key function of the PYM1-EJC interaction is to limit such EJC occupancy. Therefore, we propose that a key PYM1 function is to maintain/enhance EJC binding specificity.

Based on insights from previous studies combined with our findings that PYM1-deficient EJCs accumulate in non-canonical locations, a model emerges for PYM1’s role in maintaining canonical EJC binding specificity. By binding to RBM8A-MAGOH in a manner that precludes interaction of the heterodimer with EIF4A3(Bono et al. 2006), PYM1 relieves the inhibitory effect of RBM8A-MAGOH on the EIF4A3 ATPase cycle(Ballut et al. 2005; Nielsen et al. 2009). This action causes EJC destabilization, and release of EIF4A3 from RNA and hence EJC disassembly, as has been demonstrated earlier *in vitro* (Gehring et al. 2009). Acting in this manner, PYM1 function is analogous to that of UPF2, which stimulates UPF1 ATPase activity within NMD complexes to promote UPF1 release from RNA. UPF2 achieves this by interacting with the UPF1 auto-inhibitory cysteine-histidine rich domain to displace it from the UPF1 RecA2 domain thereby unlocking UPF1 ATPase activity, converting it from a closed to the open state, reducing its affinity for RNA and causing UPF1 release from RNA(Chakrabarti et al. 2011; Xue et al. 2023). By promoting EIF4A3 ATPase activity in a translation-independent manner on mRNAs and non-coding RNAs alike(Ghosh et al. 2014) (**Fig. 2** and **Fig. 7h**), PYM1 acts as a general destabilizer of the EJC, which otherwise maintains a very stable grip on RNA as indicated by EJC integrity for long durations and under high salt conditions in human cell extracts during procedures such as RIPiT-Seq(Singh et al. 2012; Mabin et al. 2018). In cells with compromised PYM1 activity, the EJCs could get “trapped” at canonical and non-canonical locations outside of translated regions (**Fig. 7h**). While this increased EJC stability will lead to accumulation of the complex over time including in non-canonical locations, it will also reduce EJC subunit recycling for their redeposition on canonical sites of newly spliced transcripts. At the same time, PYM1 depletion in HEK293 cells does not seem to reduce canonical EJC levels sufficiently to alter EJC-dependent cryptic splicing (**Supplementary Fig. 5b**), thereby further underscoring that PYM1-dependent EJC disassembly likely plays only a minor role in overall EJC recycling. The viability of Drosophila PYM null mutants suggests that even at the whole-organism level, the function of translation-independent EJC destabilization may be important only in specific contexts.

How PYM1 is targeted to RNA or EJCs to disassemble the complex remains to be seen. It is possible that its sequence-independent RNA binding activity could play a role in such targeting(Verma et al. 2023). Our results suggest that human PYM1’s interaction with the ribosome via its C-terminal region is unlikely to direct PYM1 to translating ribosomes for EJC disassembly. We could not directly investigate this possibility as the construct encoding PYM1 lacking its C-terminal region (PYM1^ΔC^) expressed very poorly in HEK293 cells as compared to the full length PYM1 or PYM1^ΔΝ^ constructs (**Supplementary Fig. 5g**). Therefore, the role of this PYM1 domain that bears sequence homology to EIF2A and its ability to bind to the ribosome remains unclear.

### PYM1 shapes EJC-dependent post-transcriptional gene expression

Even though PYM1 is a minor contributor to EJC disassembly, it plays an important role in EJC-mediated control of gene expression. The weak inhibition of NMD and that too only of the most robust endogenous NMD substrates (**Fig. 5**) further underscores that while PYM1 does contribute to EJC destabilization, it plays only a minor role. This NMD defect under PYM1 deficient conditions likely results from reduced canonical EJC occupancy (**Fig. 3a**), which in turn is an outcome of increased non-canonical binding or prolonged EJC lifetimes. Surprisingly, and contrary to the previous observations(Gehring et al. 2009), we did not observe any NMD defect in HEK293 cells upon elevation of PYM1 levels (**Fig. 5c**). This could be either due to insufficient overexpression of exogenous PYM1 in our experiments as compared to the past investigations(Gehring et al. 2009) or because of different PYM1 thresholds to alter EJC occupancy on endogenous versus reporter NMD targets.

Beyond NMD, we uncovered PYM1’s influence on mRNA stability in a gene architecture and membrane localization dependent manner (**Fig. 6**). The opposing changes in stability of TG+ and ER+ mRNAs are likely due to altered EJC binding on these groups of mRNAs (**Fig. 6c, d**), possibly driven by their divergent gene architectures. Such a conclusion is further supported by the emergence of gene architecture and GC content as key descriptors and predictors of differential binding of MAGOH^E117R^-EJC versus MAGOH^WT^-EJC (**Fig. 3d, e**) and altered gene expression in PYM1 depleted cells (**Fig. 6e, f**). It is possible that the EJC acts as a “reader” of gene architecture and contributes to differential mRNA localization to the ER membrane and TGs. Intriguingly, tethering of an mRNA to either EJC components or the ER membrane boosts its translation(Horste et al. 2023; Nott, Le Hir, and Moore 2004). It will be important to test if the two outcomes are due to a shared mechanism. The mechanism that alters mRNA stability in a PYM1 and gene architecture dependent manner remains to be determined. We speculate that sub-optimal canonical EJC occupancy on ER targeted transcripts alters mRNA packaging, and hence overall mRNP conformation(Singh et al. 2012; Pacheco-Fiallos et al. 2023), such that these mRNAs are exposed to mechanisms such as ER-linked altered stability(Efstathiou et al. 2022) or increased m6A-dependent instability(He et al. 2023; Uzonyi et al. 2023). It is important to note that while single-exon mRNAs show the highest increase in MAGOH^E117R^ EJC binding (**Fig. 4**), they do not exhibit any significant changes in expression (**Supplementary Fig. 6b**). Thus, the effect of increased (or decreased) EJC binding on mRNA expression levels is determined by additional transcript-dependent mechanisms.

Our work also sheds light on the possible flavivirus strategies for hijacking PYM1. We hypothesize that the flavivirus capsid protein-PYM1 interaction diverts PYM1 function away from cellular mRNPs to alter stability of host mRNAs localized at or around ER. This view is supported by congruent changes in gene expression (PTC+, TG+ and ER+ mRNAs) in cells depleted of PYM1 (**Fig. 6**), those expressing WNV capsid protein (**Fig. 7b, c**) or DENV capsid protein (**Supplementary Fig. 8d, e**), and the ones infected with DENV or ZIKV (**Fig. 7d-g and Supplementary Fig. 7f, g**). Sequestration of PYM1 away from cellular EJCs can offer flaviviruses multi-fold advantages. As flavivirus genomic RNA replicates in special compartments formed on ER membranes, disrupting PYM1 function can aid the virus in reshaping gene expression in ER proximal regions. Notably, TG+ mRNAs, which show the most robust upregulation upon PYM1 depletion and virus infection, include transcripts expressed by genes such as *RABGEF1*, *HSPA13* and *ANKRD49,* which have been identified by genetic screens as key host factors for flavivirus replication(Marceau et al. 2016; Savidis et al. 2016). Multiple studies have also shown that NMD serves as a key anti-viral mechanism(Ramage et al. 2015). Therefore, targeting PYM1 could also help the virus blunt the pathway. Furthermore, diminished translation-independent EJC disassembly and reduced EJC binding specificity in virus infected cells can promote the formation or stability of non-canonical EJCs on viral RNAs. Li *et al*. have reported that RBM8A can UV crosslink to the viral RNA which does not undergo splicing(M. Li et al. 2019). Consistently, EIF4A3 and RBM8A are among RBPs that bind to ZIKV and DENV genomic RNAs(Ooi et al. 2019). Specific mechanisms by which the EJC contributes to viral gene expression and replication are exciting questions for future investigations.

## Methods

### Plasmids and Cell lines

HEK293 cells expressing FLAG-MAGOH were created by stably transfecting HEK293 Flp-In TRex with pcDNA5 plasmid containing FLAG-MAGOH downstream of a tetracycline inducible promoter (previously reported in ref. (Singh et al. 2012)). This pcDNA5 FLAG-MAGOH plasmid was mutated by site directed mutagenesis to create FLAG-MAGOH^E117R^ and stably transfected into HEK293 cells. Briefly, 10^6^ cells were seeded into a 6 well plate and transfected the following day with 200 ng of pcDNA5 plasmid and 1.8 μg of pOG44 plasmid using Jetprime transfection reagent. 24 h later, cells were trypsinized and seeded into a 10 cm plate. After the cells had adhered to the surface, Blasticidin and Hygromycin containing media were added for selection (media was changed every 3 days). 2 weeks later, visible colonies were trypsinized, expanded and saved. A titration of tetracycline (0-625 ng/μl) was used to estimate the optimal concentration to induce near-endogenous expression of the FLAG-tagged protein.

Cells overexpressing HA-PYM1 and HA-PYM1ΔN were generated by stable transfection using PiggyBac plasmids modified from PB-TRE-EGFP-EF1a-rtTA plasmid (Addgene #104454). The promoter from this plasmid which harbored the Tet-ON system elements was swapped out for the CMV promoter from the pcDNA3.1 plasmid to achieve constitutive expression. HA-PYM1 constructs in this PiggyBac plasmid were co-transfected with a hyPBase transposase expression plasmid (described in ref. (Yi et al. 2021)) into HEK293 Flp-In TRex cells. A clonal pool of cells was isolated after selection in Puromycin containing media for ∼2 weeks.

PYM1-FLAG plasmid was generated from pcDNA3.1 N-FLAG empty vector using HiFi DNA assembly kit from NEB. WNV and DENV capsids were previously reported(M. Li et al. 2019). nLuc construct used as a control was generated by swapping the nanoluciferase sequence for the viral capsid sequence.

### siRNA transfection

For both RNA-seq and RT-qPCR experiments, 10^5^ cells were counted from a plate seeded on the previous day (actively growing cells) and reverse transfected with 10 pmols of siRNA using RNAiMAX reagent (0.8 μl of RNAiMAX and 200 μl OMEM per 10 pmol siRNA) in 12-well plates. Media was changed the following day and 72 hours later, RNA was collected by adding 500 μl Trizol reagent. For each experiment, the knockdown efficiency was calculated by western blot. siRNA sequences used: PYM1-AAACGUAACCUGAAGCGAA, EIF4A3-AGCCACCUUCAGUAUCUCA,

### Reverse transcription and quantitative PCR

Trizol extracted RNA was treated with RNase-free DNaseI (NEB) and Phenol-Chloroform extracted. 1 μg of DNase-treated total RNA was used for reverse transcription using Maxima H minus RT (Thermo fisher scientific). Reactions were set up as per manufacturer guidelines except for the amount of reverse transcriptase used (only half RTase used per unit RNA). cDNA thus obtained was diluted to the equivalent of 5 ng/μl of RNA and 3 μl was used per qPCR reaction. qPCRs were set up in 10 μl total volume using the BioRad SYBR green master mix. Each reaction was set up in triplicate and average Ct value was used for analysis. Relative fold changes of transcript isoforms were calculated using the ΔΔCt method. At least 3 biological replicates (samples transfected and collected on different days) were used for estimating errors. Primers used are listed in supplementary data file.

### Immunoprecipitation

For immunoprecipitation of FLAG-tagged proteins, stably transfected cells (induced with tetracycline the day before IP) were washed with PBS and lysed with ice-cold gentle hypotonic lysis buffer (20 mM Tris-HCl pH 7.5; 15 mM NaCl; 10 mM EDTA; 0.1 % Triton X-100; 0.1 % IGEPAL CA-630; 1x protease inhibitor cocktail) directly on the culture plate. The NaCl concentration was then raised to 150 mM, and the lysates were sonicated using a microtip at 10 % amplitude (total 6 seconds in 2 second pulses) to solubilize the chromatin fraction. Lysates were cleared by centrifugation at 15000 xg for 10 minutes at 4 °C and added to pre-washed α-FLAG M2 magnetic beads (30 μl beads per 10 cm culture plate) and nutated for 1-2 hours. Beads were then washed 8 times with isotonic wash buffer (20 mM Tris-HCl pH 7.5; 150 mM NaCl; 0.1 % IGEPAL CA-630) and immunoprecipitated proteins were affinity eluted using Isotonic wash buffer supplemented with 250 μg/ml 3X FLAG peptide. Eluted proteins were analyzed using western blot. For experiments using varying salt concentrations, cleared lysate was split into 6 tubes with α-FLAG M2 magnetic beads and increasing amounts of 5 M NaCl solution was added to lysate and then nutated for 1-2 hours.

### Puromycin incorporation assay

PYM1/Control knockdowns were performed as described above in HEK293 cells. 72 hours post transfection, cells were treated with 10 μg/ml Puromycin for 10 minutes. Cells were then harvested, and total protein concentrations were estimated using a BCA assay. Equivalent amounts of protein were loaded onto an SDS PAGE gel and after western blotting wereprobed with α-Puromycin antibody (Hybridoma bank clone PMY-2A4-S). Total protein loaded on gel was visualized using SYPRO ruby stain (Thermofisher #S12001), following standard protocols and imaged in a UV transilluminator.

### RIPiT-seq

RIPiT-seq was performed as previously described(Singh, Ricci, and Moore 2014). Eluates from the first anti-FLAG IP were subjected to second IP with anti-CASC3 antibody. For translation inhibition, cycloheximide (100 μg/ml) was added to cells 3 hours prior to cell lysis. CHX was also added to PBS used for washing as well as lysis buffer. 2 biological replicates were obtained and analyzed for each condition.

### Total RNA-seq

Knockdowns of PYM were carried out as described above. For overexpression, stably transfected cells expressing HA-tagged PYM1 were used. Knockdown efficiencies of 5 biological replicates were estimated by western blot and 3 samples with the highest knockdown efficiency were used for RNA-seq library preparation. RNA was isolated using Trizol and DNase treated, and its integrity was confirmed using high sensitivity RNA tapestation assay (RIN number > 8). 800 ng of total RNA thus isolated was ribo-depleted using RiboCop rRNA depletion kit v1.3 (Lexogen) and constructed into a sequencing library using a CORALL total RNA-Seq library prep kit (Lexogen). Manufacturer protocols were followed for both kits. Libraries were quantified on RNA TapeStation, mixed at equimolar ratio, and sequenced using the NovaSeq S4 PE150 platform (Novogene).

### RNA-seq data analysis

12nt UMIs at the start of read 1 were extracted for each read from the fastq file using UMItools(Smith, Heger, and Sudbery 2017). Adapter trimming was carried out with Cutadapt(Martin 2011) for the sequences AGATCGGAAGAGCACAC-GTCTGAACTCCAGTCAC and N{12}AGATCGGAAGAGCGTCGTGTAGGGAAAGAGTGT. Trimmed reads were aligned to the reference genome (GRCh38.p13) using the STAR aligner(Dobin et al. 2013) with Ensembl(Harrison et al. 2024) v100 annotations. Output bam files were indexed with samtools(H. Li et al. 2009) and deduplicated using UMItools (arguments --method=unique --multimappingdetection-method=NH were used). Deduplicated bam files were then sorted using samtools and converted to single-end fastq files using bedtools(Quinlan and Hall 2010). Fastq files thus generated were used as input for the pseudoalignment tool Kallisto(Bray et al. 2016) to quantify transcript-level abundances. Output files from Kallisto were imported into Rstudio using Tximport(Soneson, Love, and Robinson 2015) and used as input for DESeq2(Love, Huber, and Anders 2014) to identify differentially expressed transcripts after filtering out transcripts with transcript per million (TPM) values less than 0.1. For gene level analysis, deduplicated bam files were filtered to include only reads mapping to MANE select isoforms and lncRNAs.

Filtered bam files were then counted using HTSeq(Putri et al. 2022). Published RNA-seq data was downloaded from respective NCBI SRA projects listed in supplementary data file. Adapter trimming and PCR deduplication was performed when applicable based on the specific library orientation.

#### Analysis of NMD susceptible transcripts

A custom python script was used to annotate each transcript isoform as PTC+ or PTC-from Ensembl release 100 transcripts using their exon and 3′ UTR coordinates. Any transcript with a stop codon more than 50 nt upstream of the last exon-exon junction was defined as a PTC+ transcript. Transcript isoforms from the same gene which do not satisfy this criterion were annotated PTC-, as previously described(Yi et al. 2021). For stringent NMD targets, we used transcripts from the PTC+ set that showed an upregulation (log2FC > 0.58) and for which at least one PTC-isoform remained unchanged (log2FC between -0.58 and 0.58) upon SMG6/7co-depletion.

#### Analysis of ER associated transcripts

ER+ and TG+ gene lists were sourced from(Horste et al. 2023). For the set of genes labeled “Others”, any gene with an annotated transmembrane domain or a signal peptide was excluded.

### RIPiT-seq data analysis

A 12 nt sequence on read 5’ ends consisting of a 5 nt UMI, 5 nt identifying barcode, and a CC was removed using Cutadapt with the UMIs saved for each read for PCR deduplication down the line. miR-Cat22 adapter sequence TGGAATTCTCGGGTGCCAAGG was then removed from the 3’ end. Any reads less than 20 nt in length after trimming were discarded. Following trimming, reads were aligned with Star v2.6.0a using 24 threads to GRCh38. After alignment, reads with a mapping score less than 255 (multi-mapped reads) were removed. Next, reads were checked for overlap against hg38 annotations for miRNA, rRNA, tRNA, scaRNA, snoRNA, and snRNA using bedtools intersect(Quinlan and Hall 2010), and any reads overlapping by more than 50% were removed. Reads aligned to chrM (mitochondrial) were also counted and removed. Quantification of annotated, partly annotated and novel splice junctions detected in RIPiT-seq was done using the Qualimap package using default parameters(García-Alcalde et al. 2012).

#### Read distribution assignment

Ensembl v100 annotations of protein coding genes and lncRNAs were used for gene level quantification of EJC footprints. Fractions of reads corresponding to canonical EJC and non-canonical EJC regions were computed as follows. For last exon annotations, 100 bp were trimmed off the 5’ end. The canonical region was defined as starting 9 bp from the 3’ end of an exon and spanning 30 bp towards the 5’ end of the exon. The noncanonical region was defined to be at least 50bp long, starting 100 bp from the 5’ end of the exon and ending 150 bp from the 3’ end of the exon. Due to this definition, exons with length less than 300 bp did not have a noncanonical region in the analysis, and exons less than 100 bp were not included at all in the analysis. Read counts were then generated using featureCounts(Liao, Smyth, and Shi 2014) in a stranded manner with feature type ‘exon’, attribute type ‘gene_id’, and each read reduced to its 5’ most base. Gene level differential analyses of whole exons, and of selected exonic regions were conducted with DESeq2. Only genes with baseMean greater than 10 were used for subsequent analysis. In analyses where changing genes were grouped, either a significance threshold of padj < 0.05 or the top and bottom quartiles of log2 fold changes were used as indicated in the text.

#### Analysis of gene features

Ribosome occupancy used as a proxy for translation efficiency, was calculated from ref. (Wangen and Green 2020) by normalizing the Ribo-seq counts to RNA-seq counts for genes with a minimum of 10 reads.

Nucleocytoplasmic rank percentile was calculated from ref. (Neve et al. 2016) by ranking genes based on the nuclear ratio calculated by normalizing the nuclear RNA-seq counts to the sum of nuclear and cytoplasmic RNA-seq counts for genes with a minimum of 10 reads across both nuclear and cytoplasmic fractions. RNA half-lives were obtained from ref. (Agarwal and Kelley 2022) as a composite value of multiple independent half-life estimates in human cells (PC1).

### Machine Learning Analysis

Machine learning analysis with the goal of gene architecture feature selection was performed with four different sets of log2 foldchanges. 3 sets pertained to changes in EJC footprints for the comparisons (i) Mutant vs WT, (ii) CHX vs NT (WT), and (iii) CHX vs NT (Mutant). One set contained gene expression log2 foldchanges between PYM1 knockdown and control for MANE select genes. Gene architecture and sequence features were created from Ensembl v100 annotations. The intersection between genes for which DESeq2 was able to calculate an adjusted p-value and genes that had a value for each gene feature was used for analysis.

Genes for each respective comparison were divided into training, validation, and testing sets using an 80/10/10 % split. A deep, feed-forward, fully connected neural network was used. The network was designed with 11 layers, including an input layer with 34 neurons, multiple hidden layers with decreasing neuron counts (500, 400, 300, 200, 200, 200, 150, 100, 50), and a single output neuron. A total of 530,911 parameters were trained using a batch size of 256 and the Adam optimizer to predict log2 foldchanges from gene features using the training set. The number of training epochs and learning rate was optimized for each dataset. The model for E117R vs WT comparison was trained for 200 epochs with a learning rate of 5 × 10^-6^. Comparisons involving CHX treatment were trained for 250 epochs at a learning rate of 10^-5^, while the PYM1 knockdown model was trained for 350 epochs at a rate of 2 × 10^-6^. Model selection was conducted by measuring performance on the validation set, i.e. choosing the model which achieved the minimum loss on the validation set. Feature importance was obtained using feature permutation (importance) with the testing set to measure the increase in loss after permutation of each respective feature. The three features representing the one-hot encoded stop codon identity were permuted together. This process was repeated for 240 trained models with the percent increase in loss for each feature averaged over all models. Correlation values represent the Kendall Tau correlation coefficient between the log2 foldchanges and the feature values.

### GO term analysis

For CHX treatment, genes with differentially changing footprints identified based on *padj*<0.05 were used for GO term analysis. For MAGOH^E117R^ EJC footprints, PYM1 depletion, and viral capsid expression, top and bottom quartiles of fold changes were used for analysis. Functional enrichment was calculated in the DAVID web server(Sherman et al. 2022) and results were plotted in Rstudio.

### Data availability

The high-throughput sequencing data described in the manuscript has been deposited in NCBI short read archive and will be made available publicly upon publication.

## Acknowledgements

We acknowledge Dr. Holly Ramage for providing plasmids expressing 2X Strep-tagged WNV and DENV capsid proteins. We acknowledge the allocation of computational resources from the Ohio Supercomputer Center(Ohio Supercomputer Center 1987). This work was supported by grants from National Institutes of Health (R01-GM120209 and R35-GM149298) to G.S.

## Author contributions

Conceptualization, M.S., L.W. and G.S.; Investigation, M.S., L.W., M.L.S., R.D.P., S.M. and D.P. Writing - Original Draft, M.S., M.L.S. and G.S.; Writing - Review and Editing, M.S., L.W., M.L.S., R.D.P., S.M., R.B., and G.S.; Supervision, R.B. and G.S.

## Supplementary figure legends

**Supplementary Fig. 1. FLAG-MAGOH^E117R^ retains known EJC interactions except for PYM1.**

**a.** Crystal structure of PYM1 in complex with MAGOH-RBM8A heterodimer (left; PDB ID is indicated). Zoomed-in view on the right showing interactions of MAGOH E117 with multiple residues of PYM1.
**b.** Crystal structure of the EJC core showing that MAGOH E117 is not in close proximity to other EJC core proteins, CASC3 or UPF3B (left; PDB ID is indicated). Zoomed-in view on the right shows the distance between MAGOH E117 and the closest UPF3B residue.
**c.** Western blot showing the inducible expression of FLAG-MAGOH^E117R^ and the endogenous WT MAGOH in the stable HEK293 Flp-In cell line. Tetracycline concentrations used for 16-hour induction are indicated on the top and proteins detected are labeled on the left.
**d.** Western blot showing levels of proteins indicated on the right in total extract (input) or FLAG immunoprecipitation (IP) fractions under increasing salt concentrations for WT MAGOH (left) and its mutant counterpart (right). Salt concentrations used in IP are indicated above each lane.
**e.** Western blot showing levels of proteins indicated on the right in total extract (input) or FLAG immunoprecipitation (IP) fractions from cell lines indicated on top of each lane.

**Supplementary Fig. 2. Generation, quality control and functional analysis of the wild-type and mutant EJC RIPiT-seq libraries.**

**a**, **b**. Western blots showing levels of proteins indicated on the left in total extract (input), FLAG immunoprecipitation (FLAG-elution) and CASC3 immunoprecipitation of the FLAG immunoprecipitate (RIPiT elution) for cells expressing FLAG-MAGOH^WT^ (**a**) and FLAG-MAGOH^E117R^ (**b**).

**c.** Principal Component Analysis comparing all eight RIPiT-seq libraries generated.

**d-g**. Scatter plots comparing the gene level counts calculated for RIPiT-seq replicates for each condition: FLAG-MAGOH^WT^-EJC with no treatment [WT_NT] (**d**), FLAG-MAGOH^WT^-EJC with cycloheximide [WT_CHX] (**e**), FLAG-MAGOH ^E117R^-EJC with no treatment [E117R_NT] (**f**), and FLAG-MAGOH ^E117R^-EJC with cycloheximide [E117R_CHX] (**g**).

**h**. Density plot showing the distribution of fold-changes between FLAG-MAGOH^WT^-EJC and FLAG-MAGOH ^E117R^-EJC footprints (cyan) and between CHX versus no treatment of FLAG-MAGOH^WT^-EJC (orange).
**i.** Gene Ontology terms enriched among genes with decreased (down, on left) or increased (up, on right) EJC footprints upon CHX versus no treatment (top) and for FLAG-MAGOH^E117R^ versus FLAG-MAGOH^WT^-EJC (bottom). Dot size and color indicates the number of genes and GO term categories respectively, and x-axis represents adjusted *p*-value.

**Supplementary Fig. 3. Some EJC footprint metrics that are unaffected by disruption of EJC-PYM1 interaction and cycloheximide treatment.**

**a.** A meta-exon plot showing distribution of reads relative to the 5’ end of exons for each RIPiT-seq sample, indicated in the top left corner.
**b.** Pie charts showing the proportion of annotated, partially annotated and unannotated splice junctions observed in each RIPiT-seq library.

**c**, **d**. Cumulative distribution of fold-changes in EJC footprints in cycloheximide treated versus untreated samples for (**c**) canonical regions, non-canonical regions and last exons within transcripts and (**d**) among genes with varying number of exons.

**Supplementary Fig. 4. Features that influence and predict changes upon cycloheximide treatment of FLAG-MAGOHWT and FLAG-MAGOHE117R-containing EJC.**

**a**, **c**. Correlation of 34 gene sequence/architecture features with gene level fold-changes in the wild-type (**a**) or mutant EJC footprints (**c**) with and without CHX treatment.

**b**, **d**. Relative importance as calculated by machine learning analysis of 34 gene sequence/architecture features for the observed gene level fold-changes in the wild-type (**b**) or mutant EJC footprints (**d**) with and without CHX treatment. Error bars represent one standard deviation from the calculated mean feature importance over 30 runs.

**Supplementary Fig. 5. Effect of altered PYM1 levels on splicing, translation and NMD.**

**a.** Western blot showing PYM1 and EIF4A3 levels in HEK293 cells transfected with control or PYM1-targeting siRNA as indicated on top. Duration of siRNA treatment is above each lane. PYM1 levels normalized to EIF4A3 are shown at the bottom. The blot shown is a representative of at least 3 independent replicates.
**b.** RT-PCR products of splicing patterns of genes indicated on the right observed in HEK293 cells treated with siRNAs or CHX as indicated at the top. Genes were selected from refs (Blazquez et al. 2018; Boehm et al. 2018).
**c.** Left: SYPRO Ruby stain, Middle: anti-puromycin western blot, Bottom: western blot for proteins on the right with normalized PYM1 levels under each lane. Cell extracts were from siRNA transfected cells as indicated above each lane. On the right are pixel intensity profiles of anti-puromycin signal in siNC and siPYM1 samples, as indicated by matching colors of the text labels and profile lines.
**d.** Quantification of anti-puromycin western blot from 3 biological replicates. Error bars represent standard error calculated for the replicates.
**e.** Cumulative distribution of fold-changes in abundance of PTC-containing transcripts (red line) and isoforms without a PTC from the same genes (gray line) upon PYM1 depletion. Wilcoxon test *p*-values and transcript counts in each distribution are shown.
**f.** Boxplots showing distribution of fold-changes of transcript groups shown on x-axes upon depletion of NMD factors indicated on the top. Wilcoxon test *p*-values and distribution medians are indicated on each plot.
**g.** Western blots showing levels of HA-tagged PYM1 constructs and a control protein in HEK293 stable cell lines indicated above each lane. The PYM1^ΔC^ construct did not express well.
**h.** Western blot showing levels of proteins on the right in total extract (TE) pr FLAG-MAGOH IP from cells expressing HA-PYM1 proteins indicated above each lane.

**i**, **j**. Cumulative distribution plots as in (**e**) for HEK293 cells stably expressing PYM1 (**i**) and PYM1^ΔΝ^ (**j**).

**Supplementary Fig. 6. Effect of PYM1 depletion on functionally related gene groups and single exon genes.**

**a.** GO-term analysis of genes ranked based on altered expression upon PYM1 knockdown. The quartile of genes with lowest fold changes are labeled PYM1_down and the quartile of genes with highest fold changes are labeled PYM1_up. Dot size and color indicates the number of genes and GO term categories respectively, and x-axis represents adjusted *p*-value.
**b.** Cumulative distribution of fold-change in the abundance of single-exon genes (blue line) and multi-exon genes (grey line) upon PYM1 depletion. Wilcoxon test *p*-value comparing the two distributions is shown.

**Supplementary Fig. 7. Effects of PYM1 depletion on membrane targeted transcripts are not observed upon ER stress or NMD inhibition.**

**a.** Boxplots comparing the fold-changes of transcripts upon PYM1 versus control knockdown; in each case, transcripts were split into two classes based on annotation above each box. Wilcoxon test *p*-values and distribution medians are indicated on each plot.
**b.** Cumulative distribution plot showing the fold changes of non-membrane targeted transcripts grouped by exon count. Low (5 or fewer) and High (14 or more) exon groups represent the top and bottom quartiles of the distribution of exon counts per transcript.
**c.** Cumulative distribution plot showing the fold changes of ER+ and TG+ transcripts upon SMG6/SMG7 double depletion.

**d**, **e**. Cumulative distribution plots showing fold changes of NBAS (Longman et al., 2020) (**d**) and UPF1^LL^ (Fritz et al., 2022) (**e**) targets upon PYM1 depletion.

**f.** Same as in (**c**) upon thapsigargin treatment (Fritz et al., 2022).
**g.** Venn diagram showing overlaps between genes upregulated upon NBAS and UPF1LL depletion and those that significantly change upon PYM1 depletion.

**Supplementary Fig. 8. Effect of flaviviral capsid expression and infection on ER+, TG+ and PTC+ transcripts.**

**a.** Western blot showing levels of proteins labelled on the right in input and FLAG-IP fractions of cells expressing proteins indicated above each lane.
**b.** Western blot showing the expression levels of proteins indicated on the left in multiple replicates of HEK293 cells transfected with Strep-tagged capsid and Nluc control protein as indicated on the top. Replicates one to three were used for RNAseq library preperation.
**c.** PCA plot of RNA seq libraries indicated in the legend on the right.

**d**, **f**. Cumulative distribution of fold changes in expression of ER and TG transcripts as compared to other transcripts (**d**) and in PTC+ versus PTC-transcripts (**f**) in HEK293 cells expressing DENV capsid protein.

**e**, **g**. Cumulative distribution of fold changes in expression of ER and TG transcripts as compared to other transcripts (**e**) and in PTC+ versus PTC-transcripts (**g**) in human neuronal stem cells (hNSC) infected with ZIKV.

