## Supplementary Fig for "PYM1 limits non-canonical Exon Junction Complex occupancy in a gene architecture dependent manner to tune mRNA expression"

### Supplementary figure 1

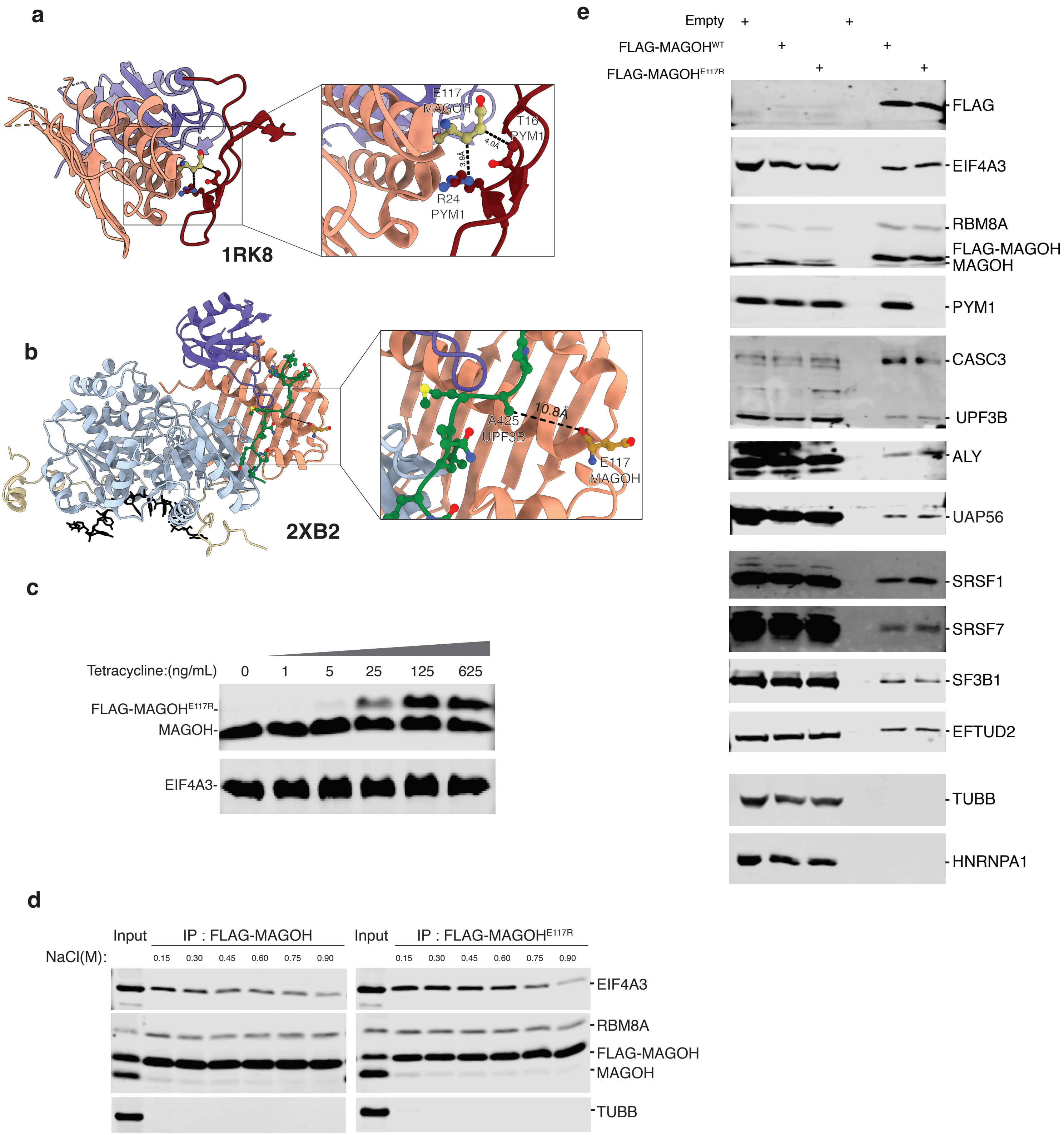

### Supplementary figure 2

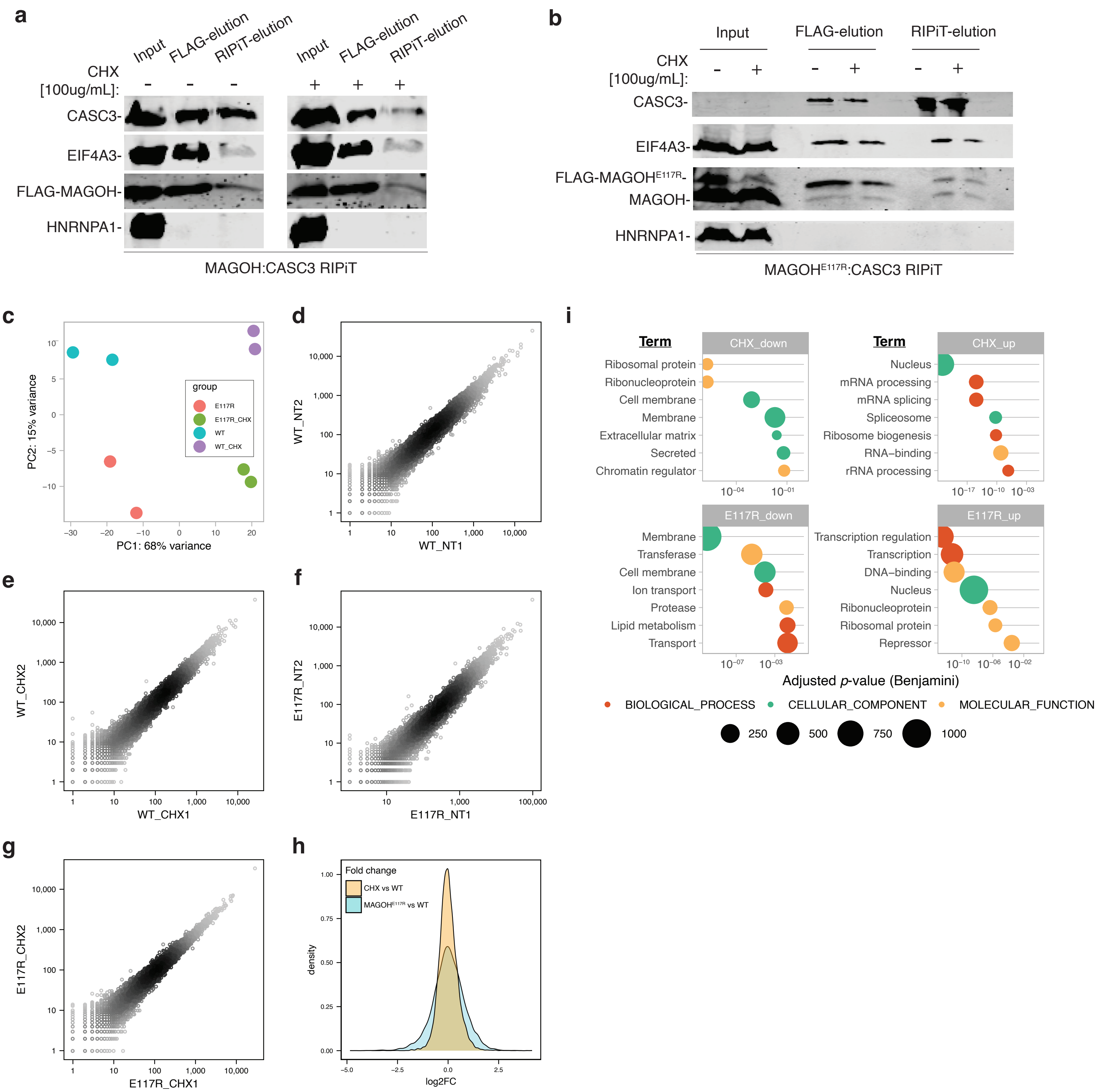

#### Supplementary figure 3

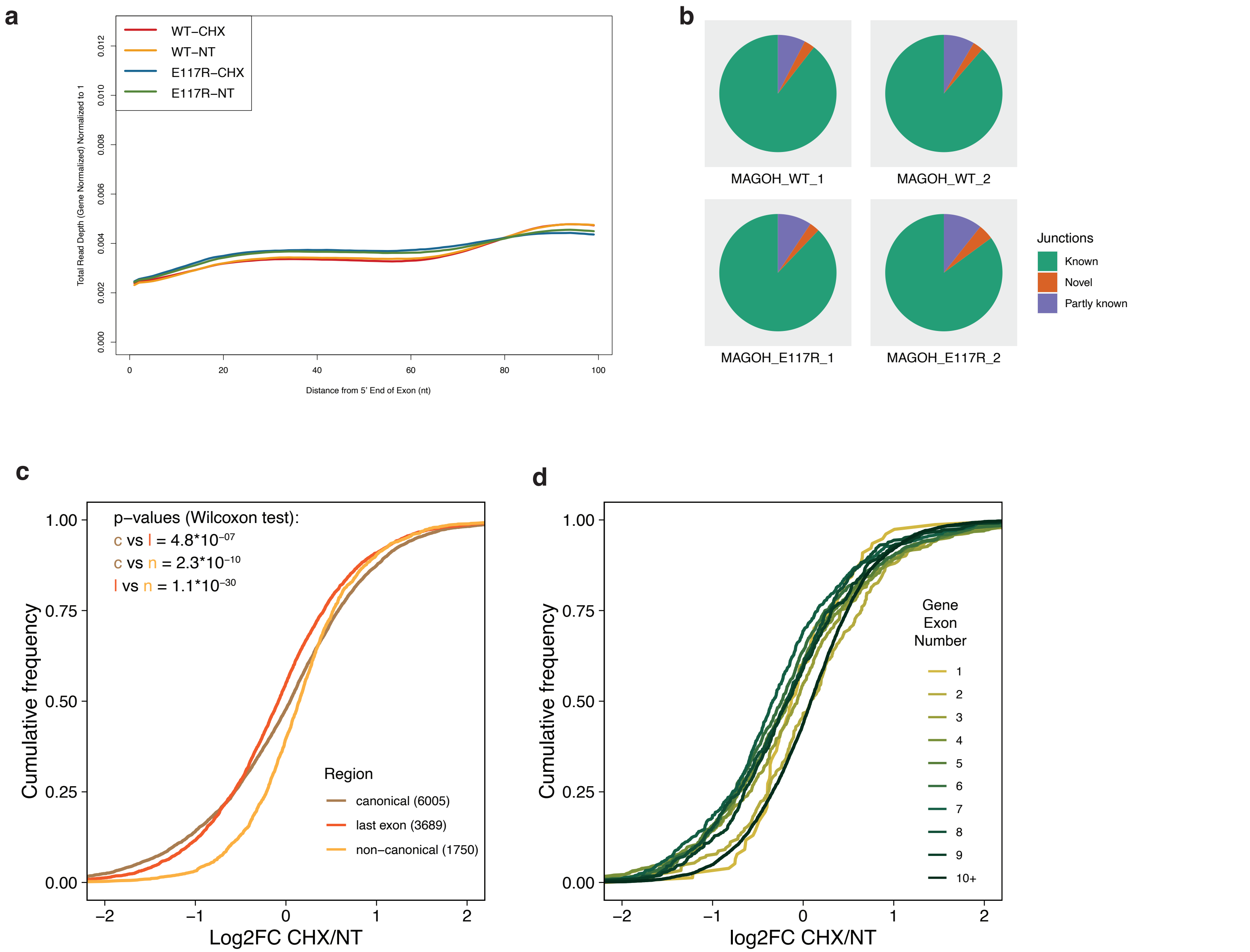

#### Supplementary figure 4

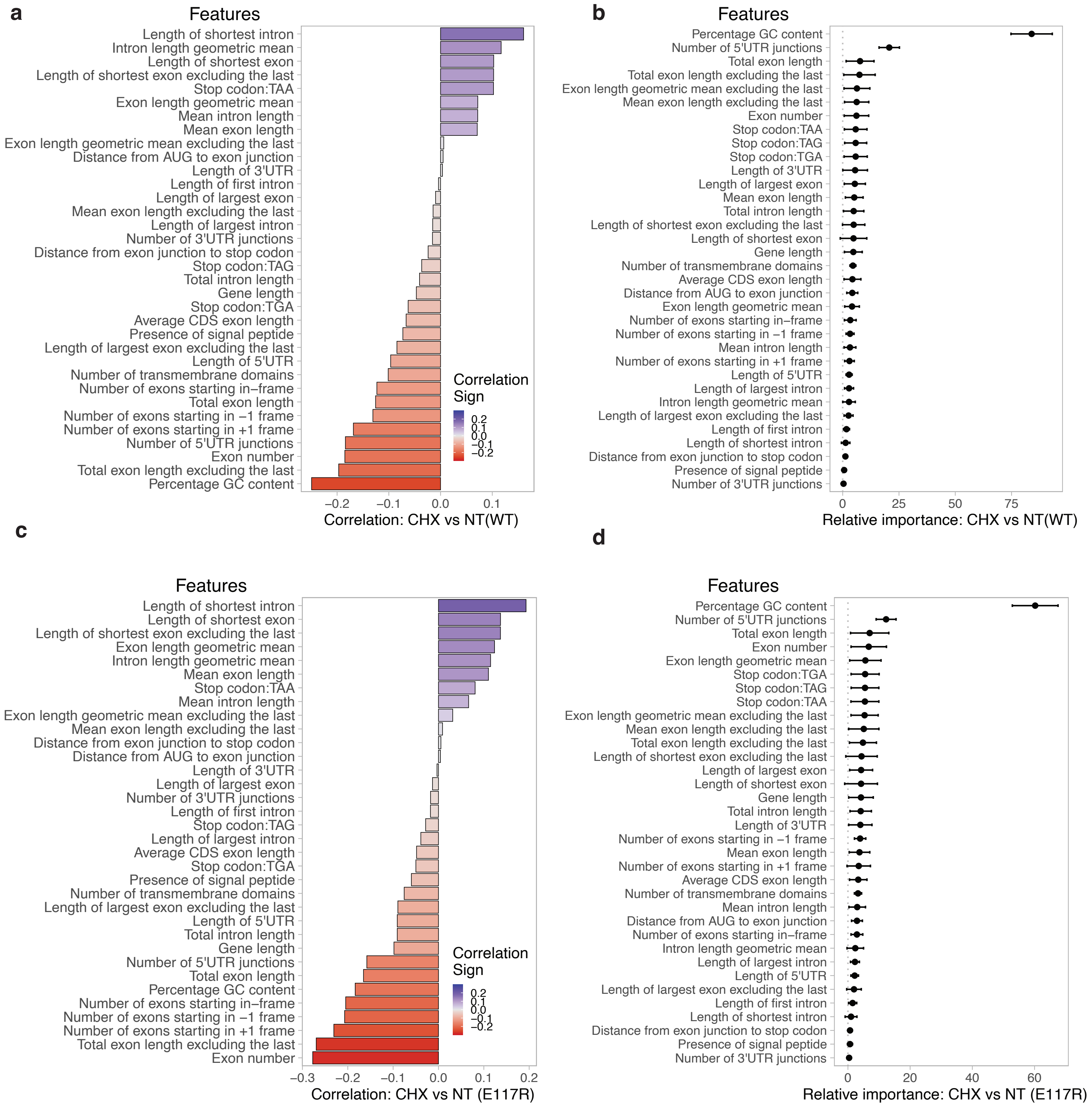

### Supplementary figure 5

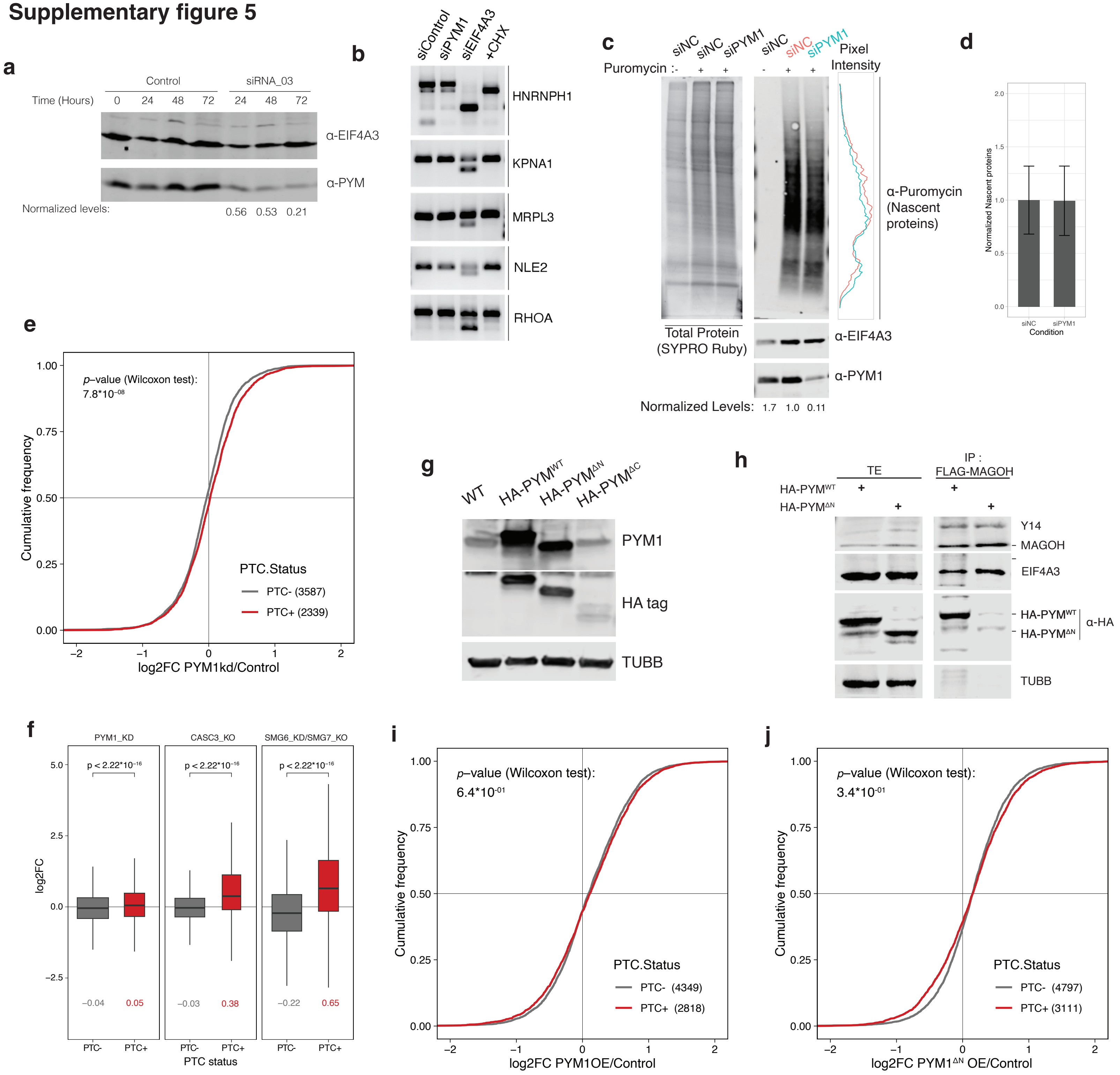

### Supplementary figure 6

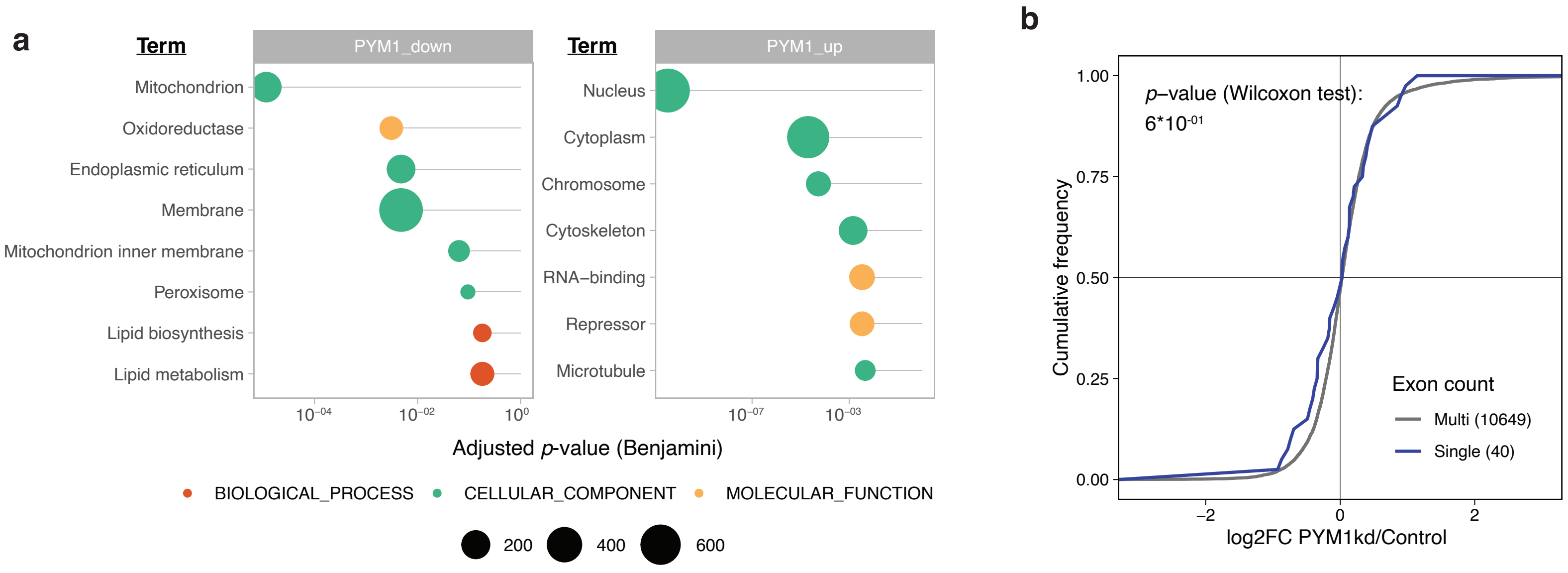

### Supplementary figure 7

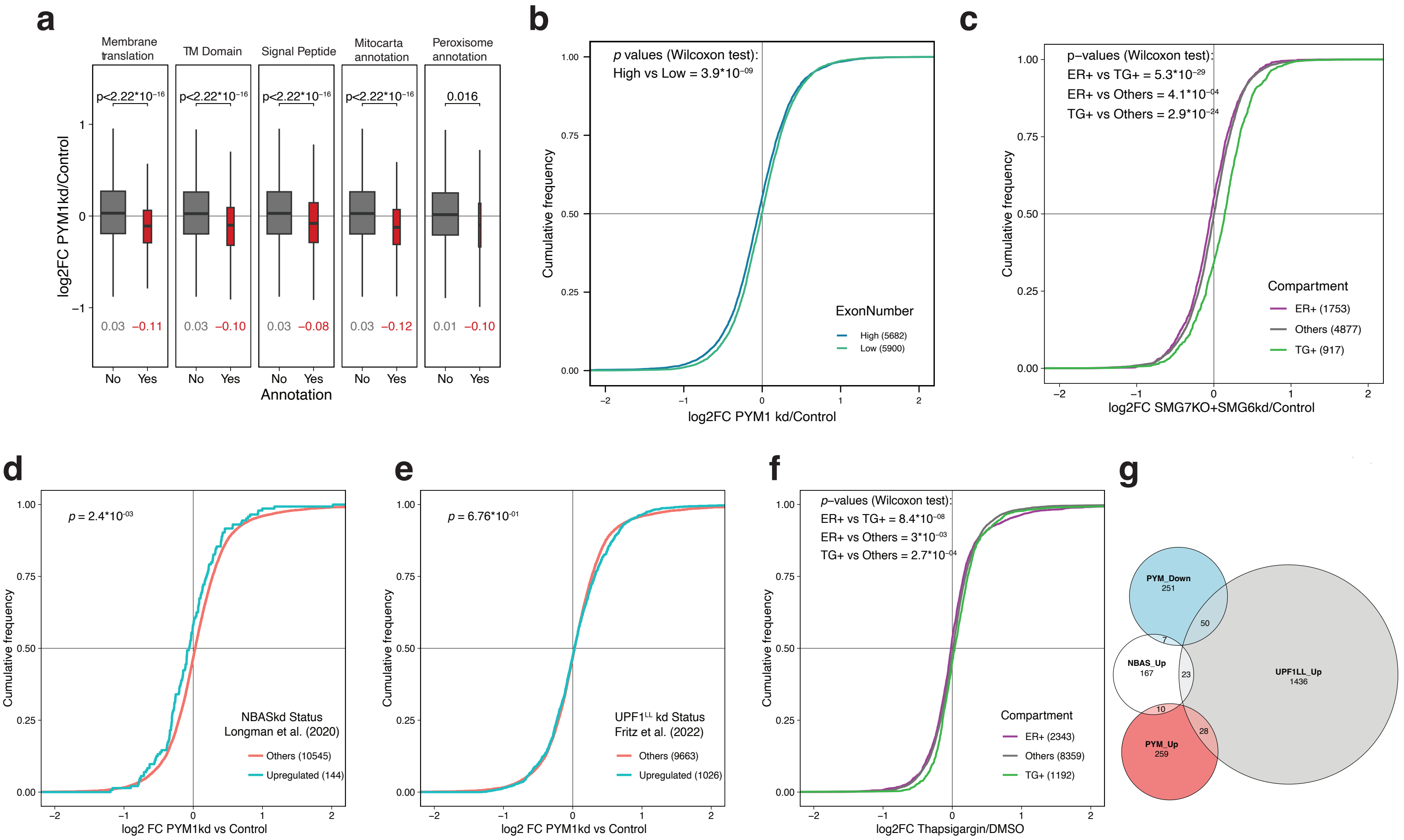

### Supplementary figure 8

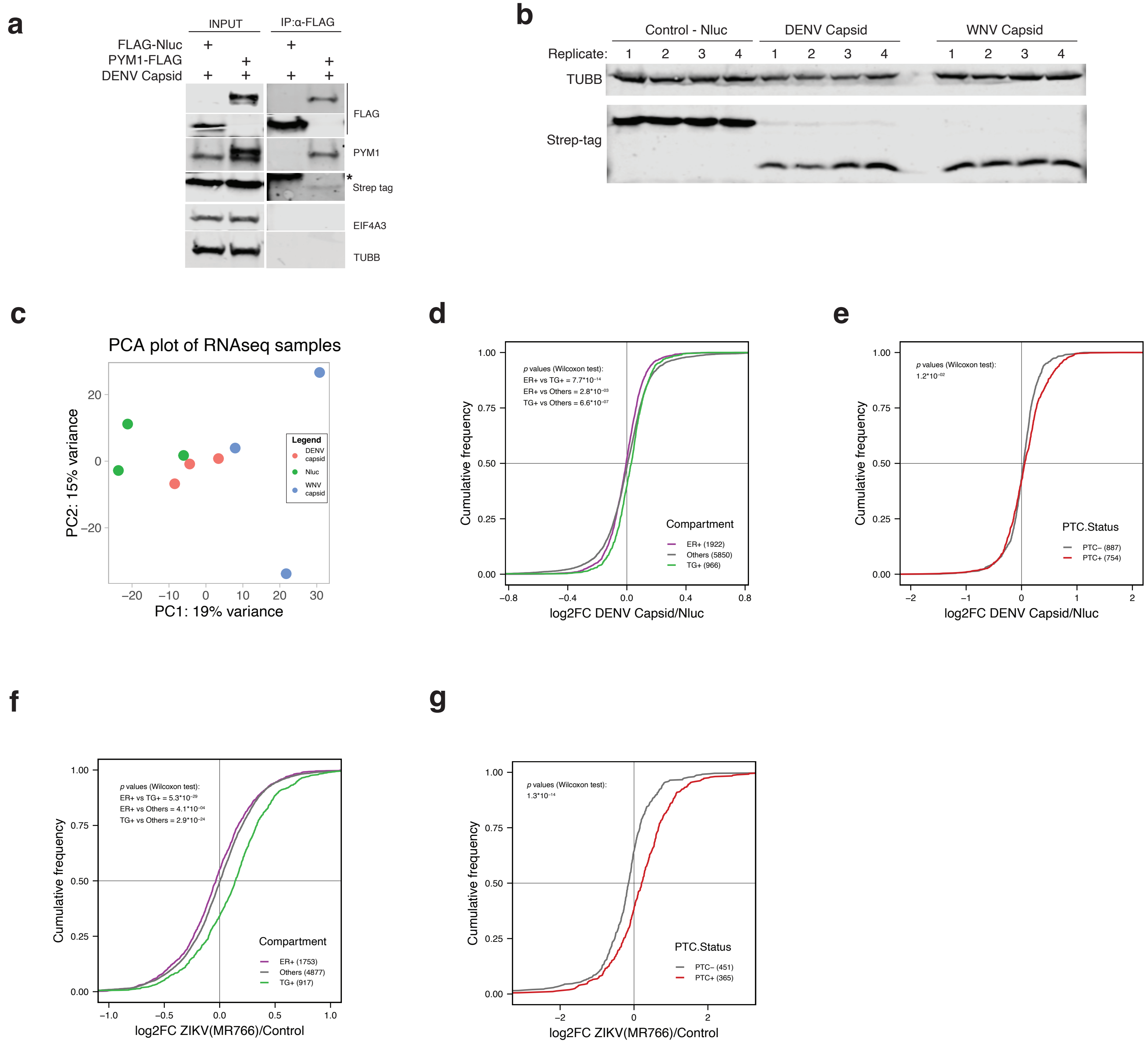
